# Meltwater runoff from the Greenland Ice Sheet reveals microbial consortia from contrasting subglacial drainage systems

**DOI:** 10.1101/2020.05.26.116566

**Authors:** Guillaume Lamarche-Gagnon, Alexandre M. Anesio, Jemma L. Wadham, Jakub D. Zarsky, Tyler J. Kohler, Elizabeth A. Bagshaw, Jon Telling, Jon R. Hawkings, Marek Stibal

## Abstract

Ice sheets overlay active and putatively widespread microbial ecosystems. An active subglacial biota has the potential to impact strongly on the (bio)geochemistry of local as well as downstream environments. Such impacts partly depend on the distribution of microbial populations, the types of habitats present beneath the ice, and their connectivity. In the ablation zone of the Greenland Ice Sheet (GrIS), supraglacial meltwaters are routed to the ice-sheet bed during the melt season, flushing out subglacial waters, sediments, and cells to proglacial environments via runoff. Here, we report on the diversity, composition, and niche differentiation of microbial assemblages exported in bulk runoff from a large (~600 km^2^) GrIS catchment. Proglacial river samples were collected over a period of subglacial drainage evolution in order to capture potential shifts in exported microbial community alongside hydrochemical transitions. We use high-resolution hydrochemical and hydrological information from the proglacial river to guide microbial (16S rRNA gene) interpretations. Core populations closely matched sequences previously isolated from other (pro)glacial environments, and phylogenetic characterisation of main OTUs alluded to a central role for subglacial iron, sulphur, and methane cycling. Whilst results indicate that bulk populations exported are likely true members of sub ice-sheet communities, we also find evidence of a supraglacial signature influencing composition of exported assemblages. Changes in assemblage structure accompanied those of major hydrological periods, with enhanced subglacial flushing coinciding with distinct shifts in microbial composition. Timing of sampling therefore matters when attempting to infer more nuanced changes in exported communities, or reveal the biogeochemical processes likely occurring in regions of the bed less influenced by surface melt. This is likely especially true when studying larger glacial systems, which experience complex hydrological changes throughout the melt-season, and that periods of extensive subglacial flushing offer opportunities to assess diversity from more isolated regions of the bed. Still, an apparent strong buffering signal from marginal zones appear to mask some of the diversity intrinsic to more remote, likely anoxic, subglacial niches, which may ultimately only be sampled via direct access to the subsurface.

## Introduction

The beds of glaciers and ice sheets contain liquid water and saturated sediments that are hosts to indigenous, active microbial communities (Skidmore *et al.*, 2000, Yde *et al.*, 2010, Christner *et al.*, 2014). Indirect observations and microcosm experiments suggest that microbial activity has an impact on both subglacial and downstream environments; e.g. by catalysing weathering reactions beneath the ice (Sharp *et al.*, 1999, Skidmore *et al.*, 2005, Mitchell *et al.*, 2013, Montross *et al.*, 2013) or via the generation and build-up of subglacial methane reserves (Stibal *et al.*, 2012, Wadham *et al.*, 2012, Burns *et al.*, 2018, Christiansen & Jørgensen, 2018, Lamarche-Gagnon *et al.*, 2019). But access to the subglacial environment remains difficult, and has often been limited to point sampling (e.g. of single marginal basal ice blocks or sediment cores) with poor temporal and spatial resolution (e.g. Stibal *et al.*, 2012, Doyle *et al.*, 2013, Montross *et al.*, 2013, Christner *et al.*, 2014).

Sampling of rivers draining land-terminating glaciers offers indirect access to the subglacial system. During the ablation season, surface meltwaters are routed to the glacier bed, flushing subglacial waters, sediments, and concomitantly microbial cells to glacial margins and proglacial landscapes. An increasing number of studies have taken advantage of such approach, broadening our understanding of (sub)glacial microbial diversity worldwide (e.g. Wilhelm *et al.*, 2013, Dieser *et al.*, 2014, Cameron *et al.*, 2017, Žárský *et al.*, 2018, Kohler *et al.*, 2020). However, glacier hydrological systems change over the course of the melt-season, which may influence the interpretations one can make from proglacial samples. Glaciers typically undergo a transition from tortuous-flow, slow and inefficient subglacial drainage during early melt (distributed system), to efficient fast-flow subglacial drainage in later months (channelised system; Davison *et al.*, 2019). Proglacial rivers are consequently sourced from waters of varying residence time beneath the ice depending on the state of the hydrological system. Timing of sampling can therefore influence, and potentially skew, interpretations if no additional information on the state of the glacier’s hydrological system is considered. Knowledge on the provenance of subglacial waters (e.g. subglacial residence time, degree of rock-water contact and weathering) also has the potential to inform on separate ecological niches present beneath the ice (Tranter et al., 2002, Tranter et al., 2005).

The influence of hydrology and hydrochemistry on proglacial microbial assemblages has previously been demonstrated. For example, Dubnick *et al.* (2017) showed that structural changes in microbial community exported from the Kiattuut Sermiat glacial catchment (southern Greenland) roughly followed changes in hydrochemistry throughout the melt season. Subtle shifts in microbial assemblages have also been linked with changes in geochemistry and water residence time in outflows of a small Alaskan glacier (Sheik *et al.*, 2015). Detailed microbial investigations of large glacial catchments are still lacking. Larger glaciers and ice-sheet margins undergo more dramatic, pronounced hydrological change throughout the melt season, draining more expansive areas, and likely export older, more remote bed material to the proglacial zone than their smaller counterparts (Wadham *et al.*, 2010, Kohler *et al.*, 2017). Consequently, proglacial rivers of larger catchments might offer a more complex picture of subglacial ecosystems provided changes in hydrological evolution is also monitored during microbial sampling.

The land-terminating Leverett Glacier (LG) drains an estimated ~600 km^2^ of subglacial catchment in the southwest sector of the GrIS. Detailed studies of its proglacial river over the last decade have shed light onto many hydrological and biogeochemical processes central to our understanding of Greenlandic glaciers and their potential impacts on downstream systems (e.g. Chandler *et al.*, 2013, Hawkings *et al.*, 2015, Hawkings *et al.*, 2017). A key aspect of LG studies has been the ability to sample during periods of increased subglacial flushing (“outburst events”), normally driven by rapid supraglacial lake drainage (hydrofracturing) to the glacier bed (Bartholomew *et al.*, 2011, Davison *et al.*, 2019). A timeseries of LG proglacial microbial assemblages, and downstream mainstem Watson River, has been reported previously (Cameron *et al.*, 2017); however, it lacked the detailed bulk meltwater hydrochemical data required to link microbial diversity to subglacial hydrology. More recent investigations have also confirmed the catchment to be a net methane source and hosting active methanotrophic populations (Lamarche-Gagnon *et al.*, 2019).

Here, we provide a detailed investigation into the microbiome of the LG catchment by combining detailed hydrological and hydrochemical information collected during the 2015 melt season (Kohler *et al.*, 2017, Hatton *et al.*, 2019, Lamarche-Gagnon *et al.*, 2019), with our previous understanding of its subglacial hydrological system (Bartholomew *et al.*, 2011, Chandler *et al.*, 2013). We test whether sampling during periods of increased hydrological flushing (outbursts) allows for more complete microbial information from isolated sections of the bed, and whether the evolution in exported assemblage structure indicates the existence of different habitats and geochemical conditions in the subglacial catchment.

## Materials and Methods

### Sampling site

Leverett Glacier hydrology and hydrochemistry have been extensively studied over the last decade (e.g. Bartholomew *et al.*, 2011, Cowton *et al.*, 2012, Chandler *et al.*, 2013, Hawkings *et al.*, 2014, Hindshaw *et al.*, 2014, Clason *et al.*, 2015, Hawkings *et al.*, 2017, Kohler *et al.*, 2017). LG is a typical example of a large land-terminating (sub)glacial catchment with processes characteristic of the western margin of the GrIS (Hawkings *et al.*, 2015, Davison *et al.*, 2019). Davison *et al.* (2019) reviews our current understanding of the hydrology of land-terminating regions of the GrIS, including schematics (notably figures 6-8 therein), which are especially relevant to conceptualise the subglacial habitats described here. Snowline distance and surface digital elevation models put the surface catchment area of LG at around 1,200 km^2^ (Palmer *et al.*, 2011; therein LG is referred to as RG), but an area of 600 km^2^ more accurately depicts the subglacial catchment described here (Cowton *et al.*, 2012). LG underlays Precambrian orthogneiss and granite, common lithology to much of Greenland (Dawes, 2009) and its proglacial river is the main source of the Akuliarusiarsuup Kuua, itself a tributary to the Watson River near the town of Kangerlussuaq. A detailed description of LG can be found in the Supplementary Information of Lamarche-Gagnon et al. (2019).

### Water sampling of molecular samples

A brief methods description of water sampling, DNA extraction, sequencing, and bioinformatics analyses can be found in Lamarche-Gagnon *et al.* (2019). Specifically, between ~ 600 – 2000 mL of LG bulk runoff was filtered through Sterivex filters (Millipore, USA) between June 07 and July 26 2015. Samples taken earlier (May 04-13) were also collected beneath river ice through boreholes and a chainsawed hole in front of the LG prior to the onset of the melt-season, and a set of samples collected on June 7^th^ from a subglacial upwelling through river ice ~ 50 m from the glacier’s terminus (see Lamarche-Gagnon *et al.* (2019) for details). The remainder of samples were collected approximately 500 m downstream from the LG portal (Fig. 1). Supplementary Table 1 details sampling volume, location and pooling of replicate samples. Waters were either collected using 60 mL plastic syringes, or a peristaltic pump (Portapump-810, Williamson Manufacturing) equipped with silicon tubing following extensive flushing of the tubing with sample water. Sterivex filters were preserved in MoBio RNA LifeGuard solution (MoBio Laboratories, USA) immediately after sampling and frozen inside a portable freezer (<-10°C) within 1 hour of collection.

**Figure 1.**
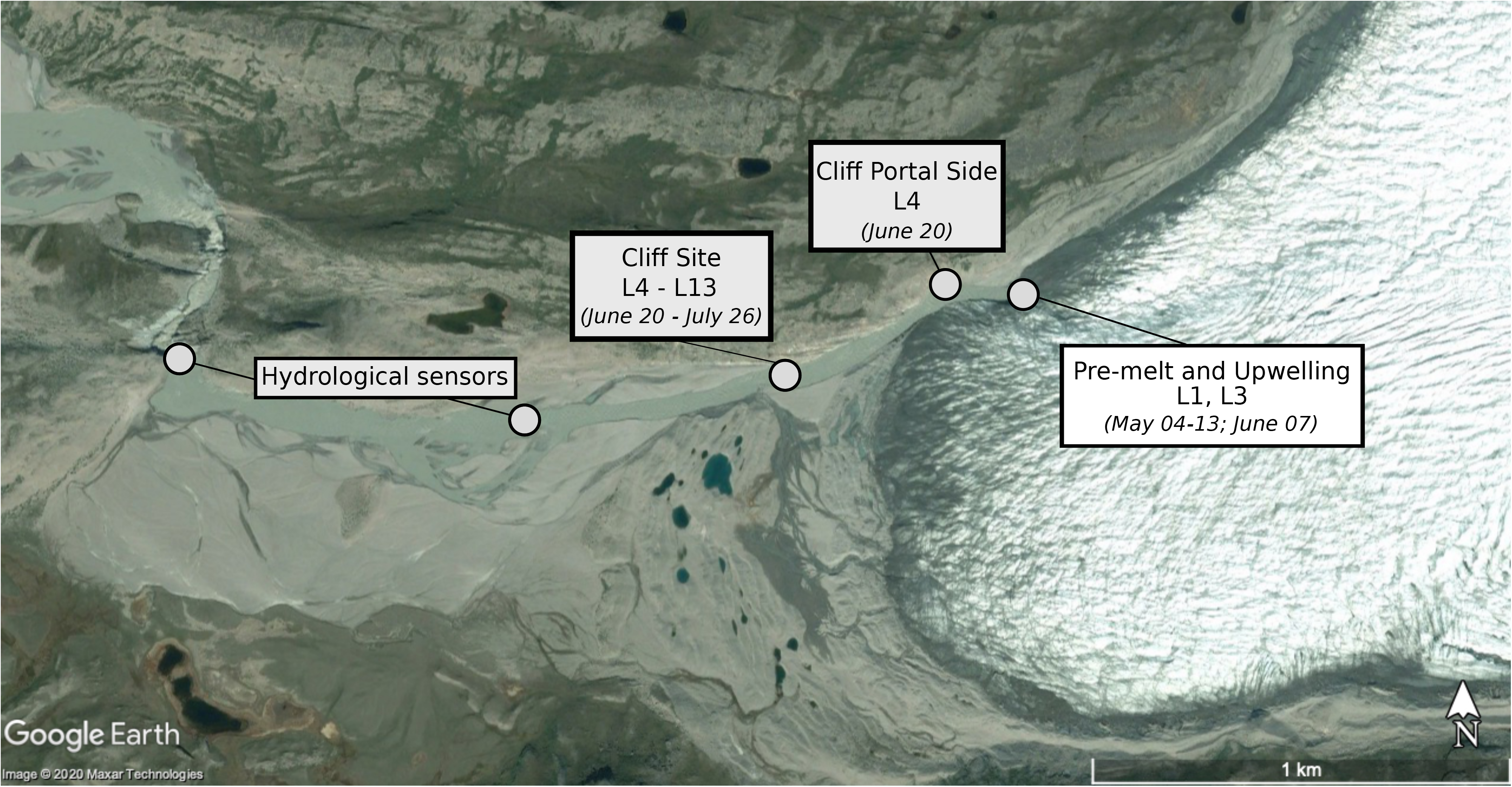
Sampling locations of proglacial river waters in front of Leverett Glacier, Southwest Greenland. Labels (L1-L13) correspond to sampling times in Fig. 2. Sampling-time details are included in Supplementary Table 1.

### Molecular analyses and initial sequence processing

DNA was extracted using the DNeasy PowerWater Sterivex kit (MoBio Laboratories, USA) following the manufacturer’s protocol. Extracted DNA samples were then pooled into triplicates based on sample date prior to sequencing, with the exception of the L1 and L3 samples (Table S1). Sequencing was performed at the Mr. DNA Molecular Research facility (Shallowater, TX, USA; http://www.mrdnalab.com/) on an Illumina MiSeq platform using the 515F/806r primer pair (Caporaso *et al.*, 2011), which targets the 16S rRNA V4 hypervariable region.

Raw sequences were analysed on the mothur platform v.1.38.1 (Schloss *et al.*, 2009) on a remote server, mostly following the mothur MiSeq standard operation procedure (Kozich *et al.*, 2013) – full details on the specific mothur commands used are provided as Supplementary Information. In short, sequences were binned into operational taxonomical units (OTUs) at a 97% sequence identity level and classified against the SILVA (v.123) database (Quast *et al.*, 2012), following quality and chimera checks. OTUs composed of two reads or less (doubletons) were removed from further analyses. Downstream analyses (e.g. beta diversity analyses) were performed on a local machine on mothur v.1.37.5. Visualisation was performed in R (version 3.5.0) (Team, 2018); the phyloseq package (McMurdie & Holmes, 2013) was also used for basic analyses. The 16S rRNA gene sequence data are available from the NCBI Sequence Read Archive (https://www.ncbi.nlm.nih.gov/sra) under BioProject PRJNA495593.

### Biodiversity analyses

Beta diversity amongst all samples was visualised using principal coordinate analyses (PCoA) and non-metric multidimensional scaling (nMDS) on an OTU similarity matrix calculated using the Bray-Curtis calculator in mothur. Prior to the generation of the similarity matrix, samples were first randomly subsampled to an equal number of reads (62,242); subsampling resulted in the exclusion of samples collected on July 1^st^ (L7s) and L8A (see Table S2) due to their lower read numbers. Samples L7s and L8A were excluded from all analyses. The significance of clustering based on the hydrological state of the LG river at the time of sampling (and amongst replicates) was tested by permutational multivariate analysis of variance (PERMANOVA) using the *adonis()* function in the R vegan package version 2.5.6 (Oksanen *et al.*, 2019). Homogeneity of dispersion, to test whether groupings have statistically (dis)similar dispersions, was conducted using the *betadisper()* function in vegan.

In order to gain further information on OTUs influencing the ordination clusterings above, the 20 most influential OTUs (by p-value) were extracted from both the PCoA and nMDS using the *‘corr.axes’* command in mothur applying the spearman method to calculate correlation coefficients. Changes in relative abundance of these OTUs were visualised as a heatmap using the *heatmap.2()* function of the *gplots* package (Warnes *et al.*, 2019). Dendrogram organisation was produced using the Ward.2 method as it resulted in clusters more intuitive to interpretations.

### Phylogenetic tree

Sequence alignment and phylogenetic tree generation were performed in MEGA v.7.0.26. Sequences were first aligned by ClustalW using default settings. Aligned sequences were then trimmed to the region of interest (i.e. region spanning the 515 to 806 region of the 16S rRNA) and alignments visualised on maximum parsimony trees using 1000 bootstrap using the Jukes-Cantor method.

### Experimental design based on hydrological and hydrochemical evolution of the LG drainage system

To interpret changes in microbial community structure within the context of their subglacial sources of export and ecological niche transitions, we also include hydrological and hydrochemical data from LG runoff. Figure 2 puts microbiological sampling in the context of the “hydrological state” of the LG system at the time of sampling based on our understanding of the LG drainage system during the 2015 melt-season (Kohler *et al.*, 2017, Hatton *et al.*, 2019, Lamarche-Gagnon *et al.*, 2019). Here, we also underline a few additional “hydrological states” that characterised the 2015 melt-season at LG, to better constrain the potential sources of exported microbial assemblages. Broadly, we follow the same hydrochemical interpretation from Hatton *et al.* (2019) and Dubnick *et al.* (2017), whereby increases in divalent (Ca^2+^, Mg^2+^) to monovalent (Na^+^, K^+^) cation ratios (D:M) are interpreted to reflect a larger contribution of carbonate over silicate mineral weathering reactions to the cation load, respectively hypothesised to derive from subglacial waters of shorter to longer residence times subglacially (see below).

**Figure 2.**
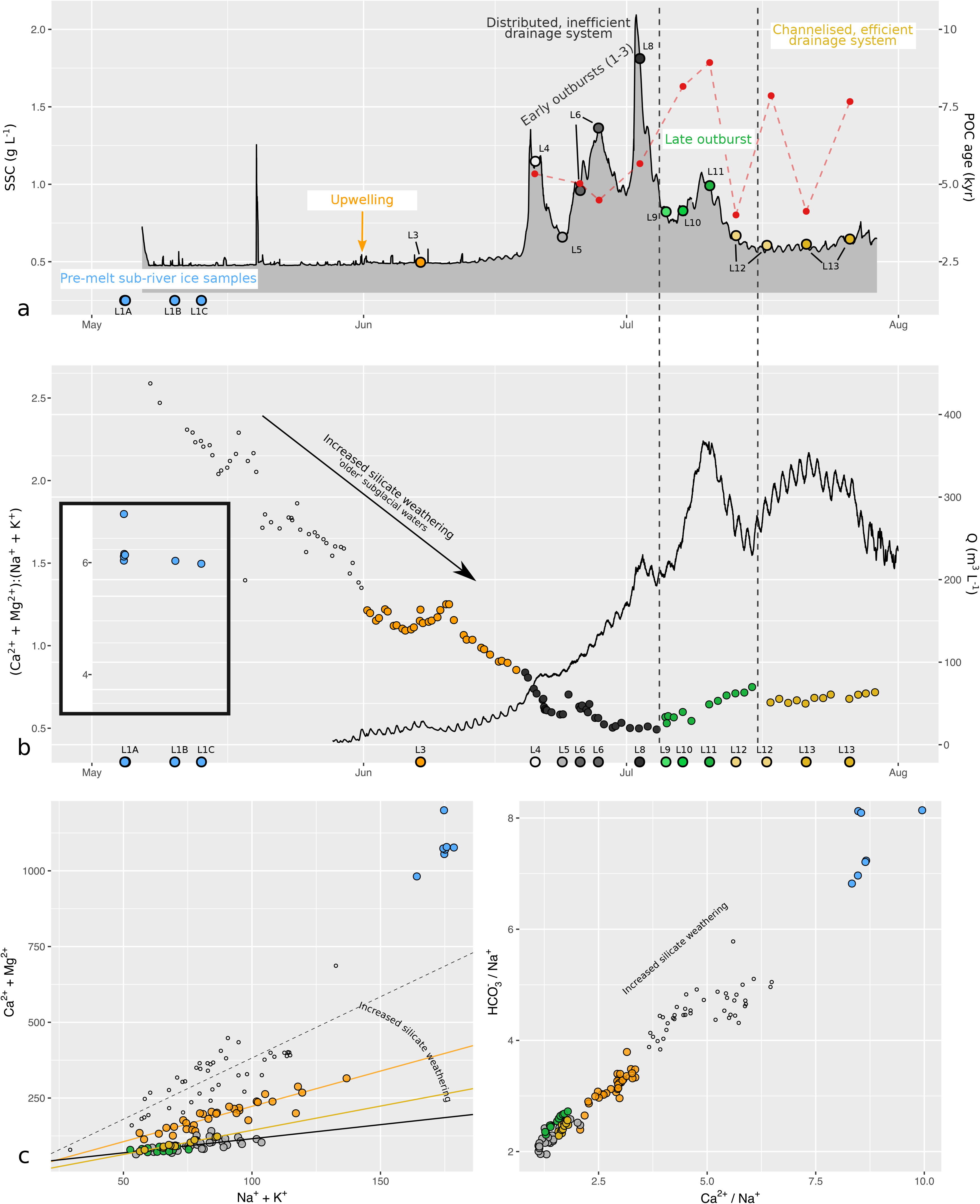
Hydrogeochemical evolution of LG proglacial river and sampling details – (**a**) Timeseries of suspended sediment concentration (SSC; black) and inferred age of suspended-sediment-associated particulate organic carbon (POC) based on ^14^C dating in red (data from Kohler *et al.*, 2017). Bordered coloured points indicate sampling time of waters used for DNA extraction overlaid onto the SSC timeseries; L1 samples were collected beneath river ice and no SSC data is available for those samples (see methods). The four abrupt increases in SSC between June 19^th^ and July 15^th^ correspond to outburst events (Lamarche-Gagnon *et al.* (2019); Kohler *et al.* (2017)). Approximative hydrological states of the LG drainage system are highlighted. (**b**) Discharge timeseries (black line) and ratios of major divalent-monovalent cations (D:M; points) of LG runoff (Hatton *et al.*, 2019). Colour of points reflect LG hydrological states; inset shows D:M of sub-river ice waters – note the difference in scale on the y-axis (data provided as Supplementary Information). Sampling time of waters used for DNA extraction are also highlighted as per **a** on the x-axis. (**c**) Major ion relationships diagrams (D:M and Na^+^-normalised molar mixing ratios of Ca^2+^ and HCO_3^-^_ of LG water samples alluding to increased/decreased silicate weathering; samples and colours are the same as in **b**. Lines on the left panel are linear regressions for samples taken before the emergence of the upwelling (dashed line), following the appearance of the upwelling but prior to the first outburst (orange line), during the outburst period (black line) and following the last outburst (amber line). See Hatton *et al.* (2019) for more detailed interpretations.

The evolution of the subglacial drainage system at LG and its connectivity to the proglacial river can be inferred from changes and evolution in discharge, turbidity, and major ion chemistry, summarised in Figure 2. Early samples (L1) were collected from relatively stagnant or slow moving waters beneath the river ice in early May; L3 samples were collected following the emergence of a subglacial upwelling through river ice (~ June 01), which was accompanied by a switch in the relationship between major divalent and monovalent cations (Fig. 2c). The emergence of the upwelling also resulted in a rise in dissolved methane concentrations at the sampling site, further indicative of proglacial connectivity to the subglacial hydrological system of the marginal zone, driven by increased melt (Lamarche-Gagnon *et al.*, 2019).

The series of four pulses in SSC between June 19 and July 15 reflect the rapid drainage of supraglacial waters to the base of the glacier (Bartholomew *et al.*, 2011, Lamarche-Gagnon *et al.*, 2019). This large input of dilute meltwater mechanically disrupts the glacier’s bed, flushing out waters with high sediment loads from a distributed hydrological system (Bartholomew *et al.*, 2011). At LG, these outburst waters bear a chemical weathering signal characterised by increased silicate mineral dissolution compared to carbonate mineral dissolution, illustrated by a decrease in the D:M ratio, and reflective of longer residence time at the glacier’s bed (Fig. 2b; Hatton *et al.*, 2019). The hydrochemical evolution from decreasing to increasing D:M ratios following the third outburst event (~ July 5^th^) may reflect a small relative increase in carbonate weathering signal from carbonation and hydrolysis of trace carbonates in an increasingly efficient drainage system, following the evacuation of long residence-time, over-winter stored waters (and sediments), or “mechanically renewed” by active bedrock comminution, in an expanded drainage system experiencing longer contact with fully oxygenated, highly turbulent, surface meltwaters (Brown, 2002). The evacuation of subglacial material from more remote sources can also be inferred by the export of suspended sediments bearing an older particulate organic carbon (POC) content during this period (Figure 2a; Kohler *et al.*, 2017). Lastly, the hydrological period following the last outburst at LG (after July 15) is reflective of a fully expanded, efficient and channelized drainage system (Hatton *et al.*, 2019).

Based on changes in runoff hydrochemistry, flow regime, and SSC, we can therefore separate the microbiological sampling schedule into five time periods. We herein define these different stages as “pre-melt” (L1 samples); “upwelling” (L3 samples, June 1-18); and “outburst”. The latter is sub-divided into two: “early outburst” spanning the first 3 SSC pulses (June 19 – July 3; L4 to L8), and “late outburst” (L9 to L11 samples); and “post-outburst”, or channelised, (July 15^th^ onward; L12-L13) periods (Fig. 2).

### Hydrological and hydrochemical metadata

Methods for determination of suspended sediment concentrations (SSC) and discharge (Q) measurements can be found in Kohler *et al.* (2017), as well as Lamarche-Gagnon *et al.* (2019); processed sensor data is also available in Lamarche-Gagnon *et al.* (2019). In brief, turbidity (Partech C sensor) was converted to SSC by calibration against manual samples and discharge from pressure transducers (Druck and Hobo) and a mobile water depth sensor (Campbell Scientific SR50A) either deployed ~ 1.6 km (turbidity), or fixed in a bedrock section about 2 km, downstream from the glacier’s terminus. Pressure/stage measurements were converted to a stage-discharge rating curve generated from calibration against repeated rhodamine dye injections over the full range of river stages during the melt season, as in Bartholomew *et al.* (2011).^14^C-POC data was taken from Kohler *et al.* (2017). Methods and data regarding monovalent and divalent cation concentrations for the proglacial river can be found in Hatton *et al.* (2019); major ion data for borehole waters are included here as supplementary information (Table S3). Borehole waters were collected using the peristaltic pump as for the collection of microbial samples. Storage and analyses of those samples were identical to that described in Hatton *et al.* (2019); analyses were also performed at the same time of the LG samples presented there. Briefly, water samples for major ion analyses were filtered through 0.45 μm cellulose nitrate filters (Whatman) mounted on portable filtration stack units (Nalgene) and stored refrigerated in the dark until analyses. Major ion concentrations were determined by ion chromatography (Thermo Scientific Dionex capillary ICS-5000) (Hawkings *et al.*, 2015).

Dissolved oxygen, pH, and EC concentrations in borehole waters were measured using an Aanderaa Optode 3830 sensor, Honeywell Durafet pH sensor and Campbell Scientific 547, respectively (Table S3).

## Results

### Community composition

Overall, the bulk of exported microbial communities retained a stable make-up throughout the melt-season, with Proteobacteria, Bacteroidetes, Actinobateria, Chloroflexi, Acidobacteria, and Verrucomicrobia constituting the major phyla in all samples (Fig 3.a). At the order level, all samples were dominated by Burkholderiales, with a relatively stable composition of Nitrosomonadales and Sphingobacteriales (Fig. 3b). However, some important temporal differences were observed, such as a relatively large proportion of Methylococcales, and very low abundance of Flavobacteriales, present in pre-melt and late-season samples (i.e. L1 and L13), the near absence of Xanthomonadales before the melt season (L1), and a larger proportion of Anaerolineales sequences detected during the outburst period (Fig. 3b).

**Figure 3.**
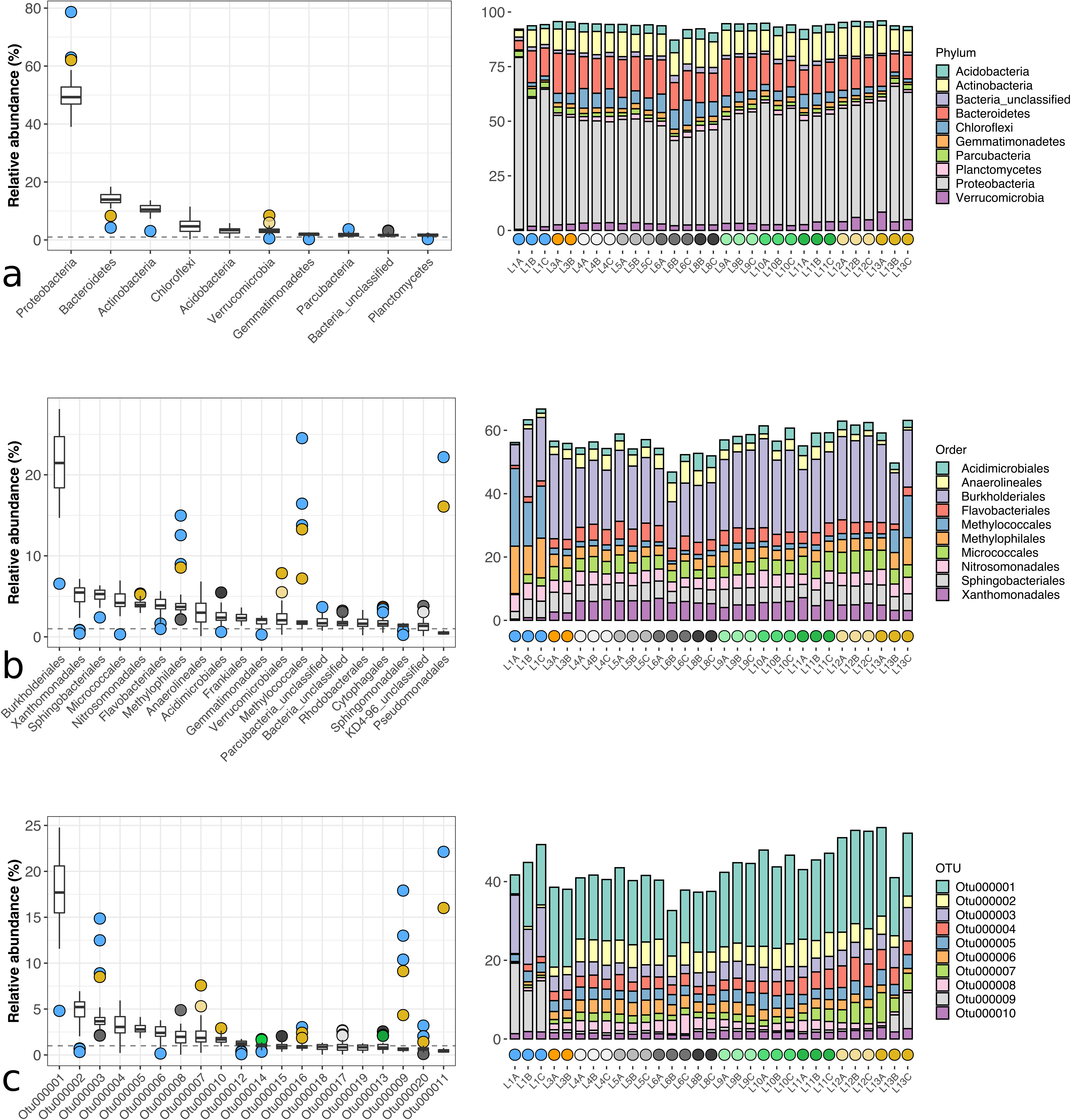
Relative abundance of major phyla (**a**), orders (**b**) and OTUs (>97% sequence similarity; **c**) in LG microbial assemblages. Data is shown as both box plot (left) and bar plot (right); only the top 10 taxa are shown in each bar plot whereas the top 20 orders (**b**) and OTUs (**c**) are shown in the box plots. The box mid-lines represent medians; the interquartile range (IQR) is represented by the lower and upper box boundaries, which denote the 25th and 75th percentiles, respectively; whiskers indicate confidence intervals 1.5 times the IQR, and points are outliers. Colour of points correspond to sampling time – same colour scheme as Fig. 2 is used. The dashed lines in the box plots mark 1 % relative abundance.

A similar pattern is retained when focusing on major OTUs (Fig. 3c, Table 1). The dominance of Burkholderiales is reflected by the most abundant OTU across all samples (OTU 1). Differences between pre-melt and post-outburst samples and the rest of the season are also reflected with the relative abundance of specific OTUs. That is, a much larger relative abundance in OTU 2 (Xanthomonadales), 6 (Flavobacteriales), and 8 (Anaerolineales) is observed for samples collected during the outburst periods, whereas OTU 3 (Methylophilales), 9 and 16 (Methylococcales), and 11 (Pseudomonadales) were most abundant in the pre-melt and latest samples (L1 and L13; Fig. 3c). OTU 7 (Verrucomirobiales) is also markedly more abundant during the post-outburst period (L12-13). Archaea only represented a minority of sequences, with the most abundant archaeal OTUs accounting for less than 0.1% sequences in most samples (Fig. S1). Of note is OTU 137, made up of Methanosarcinales sequences related ANME-2d anaerobic methane oxidising archaea, which also exhibited a marked increase in relative abundance in pre-melt and post-outburst samples (Fig. S1-2).

**Table 1.**
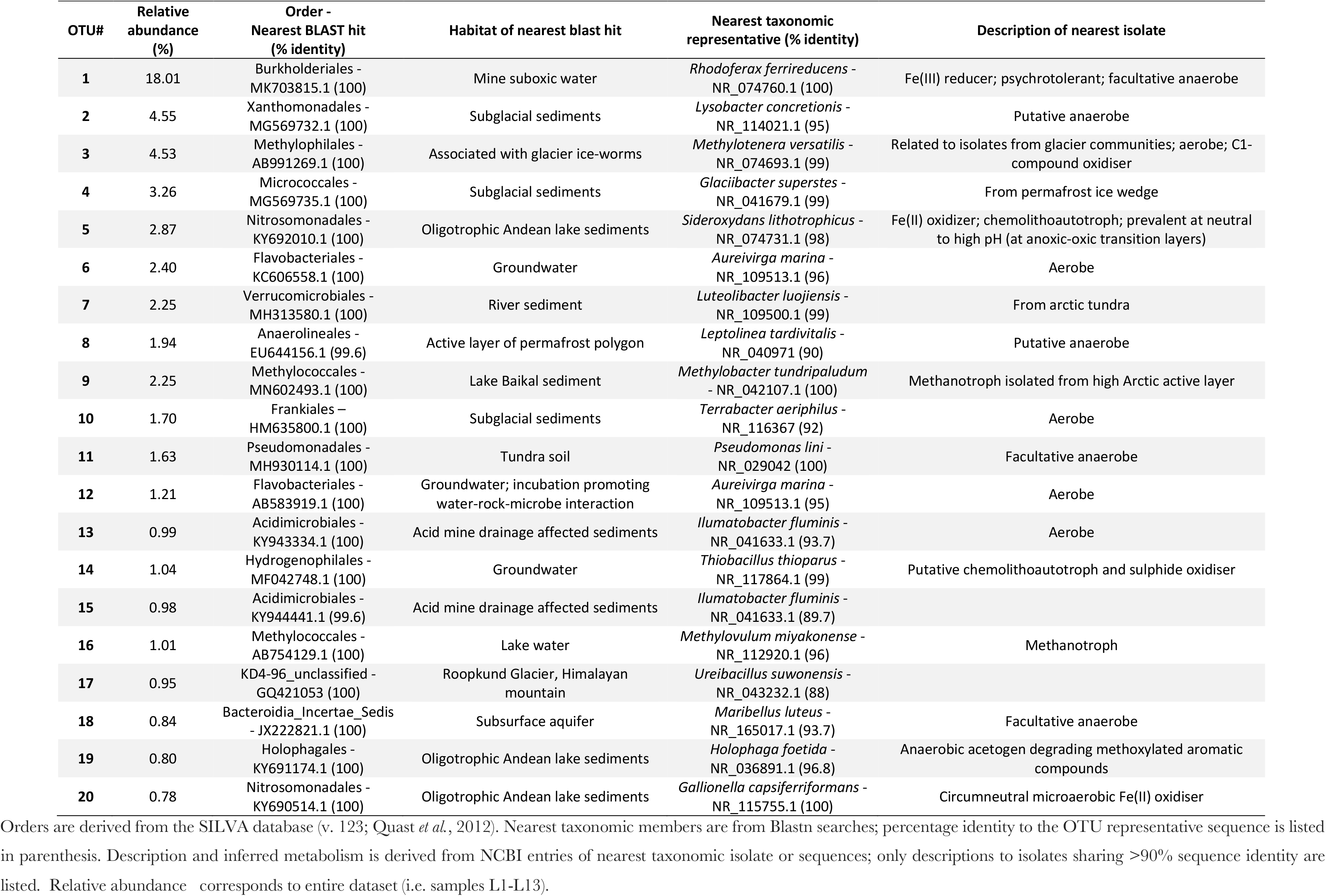
Taxonomic and inferred metabolic description of LG dominant OTU representative sequences.

### Temporal evolution of exported microbial assemblages

Whilst focusing on the most abundant populations did not allude to major community changes during the sampling period, whole-community comparisons, however, did highlight specific patterns in community assemblages and reveal a temporal evolution in exported microbial community structure that closely followed changes in hydrochemical states of exported waters. The most distinct microbial communities are observed for samples collected prior to the onset of the melt-season (L1) and the very last set of samples collected during the period of efficient (channelised) subglacial drainage (L13) (Fig. 3b, c). When communities are visualised through ordinations of Bray-Curtis dissimilarities, L1 and L13 samples indeed clearly cluster separately from those exported during the upwelling and outburst periods (L3-L11) (Fig 4a, b). Such changes are mostly illustrated by shifts along the vertical axes of the PCoA and nMDS plots, which depict the highest level of variation amongst communities. Although more subtle, a clear evolution in community structure can also be seen between samples collected following the onset of the melt-season (L3-L11), characterised by shifts along the x axes of Figure 4a, b. Differences between samples are more apparent on the nMDS plot (Fig. 4b). On the PCoA, “outburst communities” are better grouped temporally, with samples flushed out during the period of the first three outbursts (~ June 19 to July 5) forming a separate cluster to the last one (L10-L11; ~ July 6-13; Fig. 4a).

**Figure 4.**
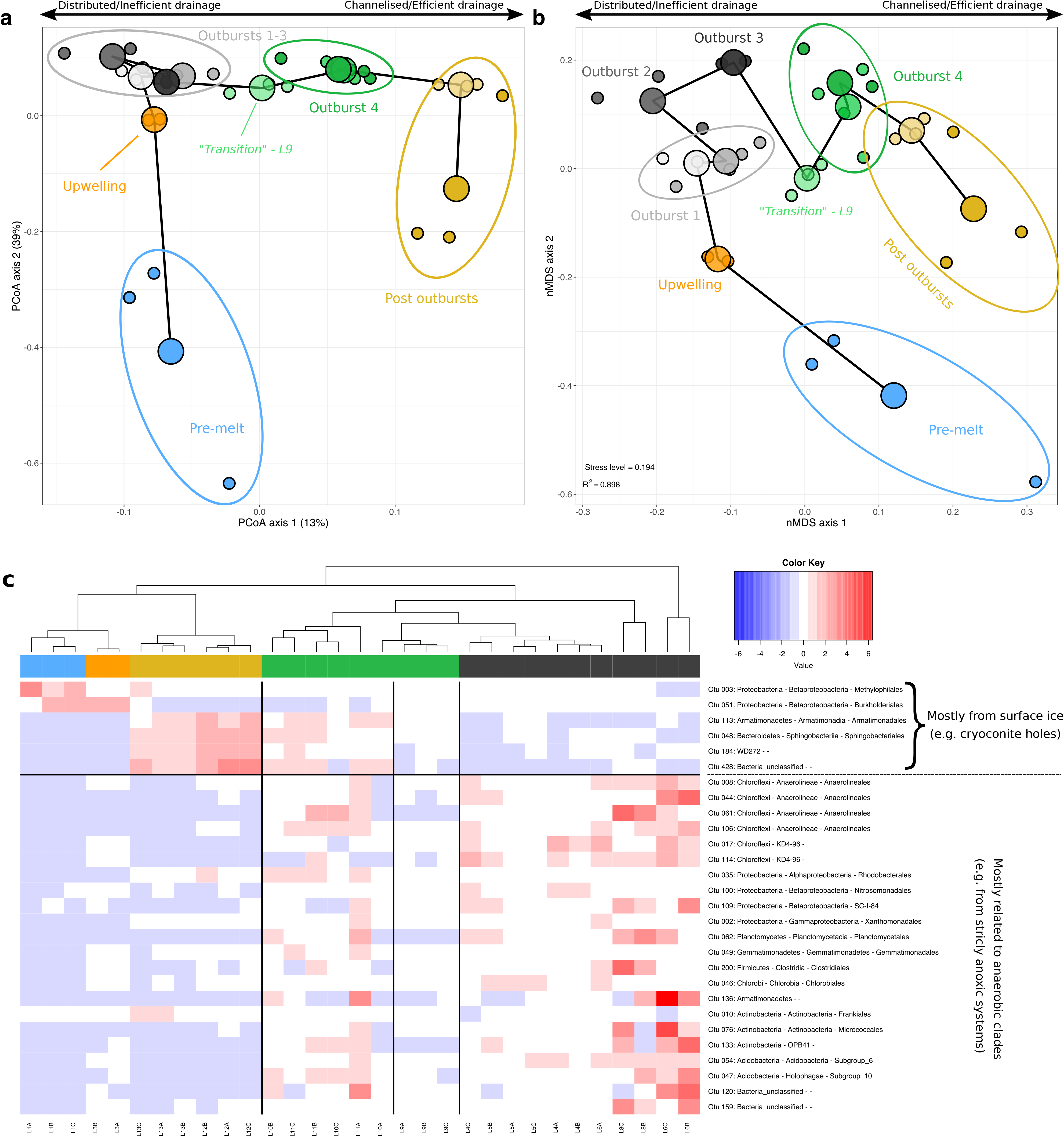
(**a**) Principal component analysis (PCoA) and (**b**) non-metric dimensional scaling (nMDS) projections of Bray-Curtis dissimilarity matrix on LG communities. Small dots represent replicate samples and large ones are averages. Colours and groupings are the same as in Fig. 2. The thick black line links average points by sampling time. On the PCoA, axes indicate explained level of variation; the nMDS stress level is 0.19, R^2^ = 0.90. Note the difference in scale between axis 1 and 2 for both ordination plots. Clusters represent communities grouped by “hydrological states” as per Fig. 2, with clusters having significantly different centroids: Permanova for clusters highlighted on the PCoA (R^2^ = 0.62, Pseudo-F = 8.21, p < 0.001); permanova for clusters highlighted on the nMDS (R^2^ = 0.72, Pseudo-F = 7.24, p < 0.001). Dispersion of homogeneity tests show no significant difference in dispersion between clusters: pseudo F = 1.00, p > 0.1 for the PCoA; pseudo F = 1.23, p > 0.1 for the nMDS. L9 samples (light green) are highlighted as “transition” between earlier outbursts and outburst 4. Duoble balck arrows for the x axis reflect the approximate spectrum of the hydrological state at the time of sampling; does not apply to the pre-melt (blue) samples. (**c**) Heatmap visualisation of the top 20 OTUs most influencing clustering on the ordination plots above (28 OTUs total; 20 shared and 8 unique to the PCoA and nMDS ordinations). Colours (blue to red) show abundance of an OTU relative to the average amongst all samples for that OTU; cooler colours indicate lower than average and warmer higher. (for relative abundance of that OTU per sample (i.e. relative to other OTUs), see Table 2). Colours of samples reflect main hydrological states as per Fig. 2. Taxonomic information down to the order level (separated by dashes) for each OTU is indicated, when available.

The “transition” in community structure observed between early and late outburst clusters coincided with a change in hydrochemical regime, highlighted by a small change in major divalent to monovalent cation relationship and age transition of exported POC (Fig. 2), likely reflective of the flushing of subglacial sediments and waters with difference residence times (Kohler *et al.*, 2017). This change in microbial assemblages between the first 3 outbursts (samples L4-L8) and the last one (samples L10s-L11s) is further illustrated by the community structure of L9 samples, collected during this outburst transition period, which roughly centre both PCoA and nMDS plots (Figure 4 a, b). It should be noted that L12 samples were homogenised from waters collected during two distinct hydrological stages; both L12 and L13 samples were also grouped from waters and sediments of different POC age, which may have resulted in dampening observed microbial changes that may have occurred during this later time period (Fig. 2).

Isolating the OTUs that most influenced ordination clustering provides further information on the potential hydrological mechanisms responsible for the restructuration of proglacial assemblages during the melt-season (Fig. 4c). Broadly, earlier and later communities appear most influenced by populations with a potential supraglacial origin, whereas a higher proportion of sequences related to clades typically associated with more hypoxic/anoxic environments appear over-represented in outburst communities (Fig. 4c, Table 2). Again, the apparent ‘transition nature’ of L9 samples is apparent by no relative increase in any of these OTUs.

**Table 2.**
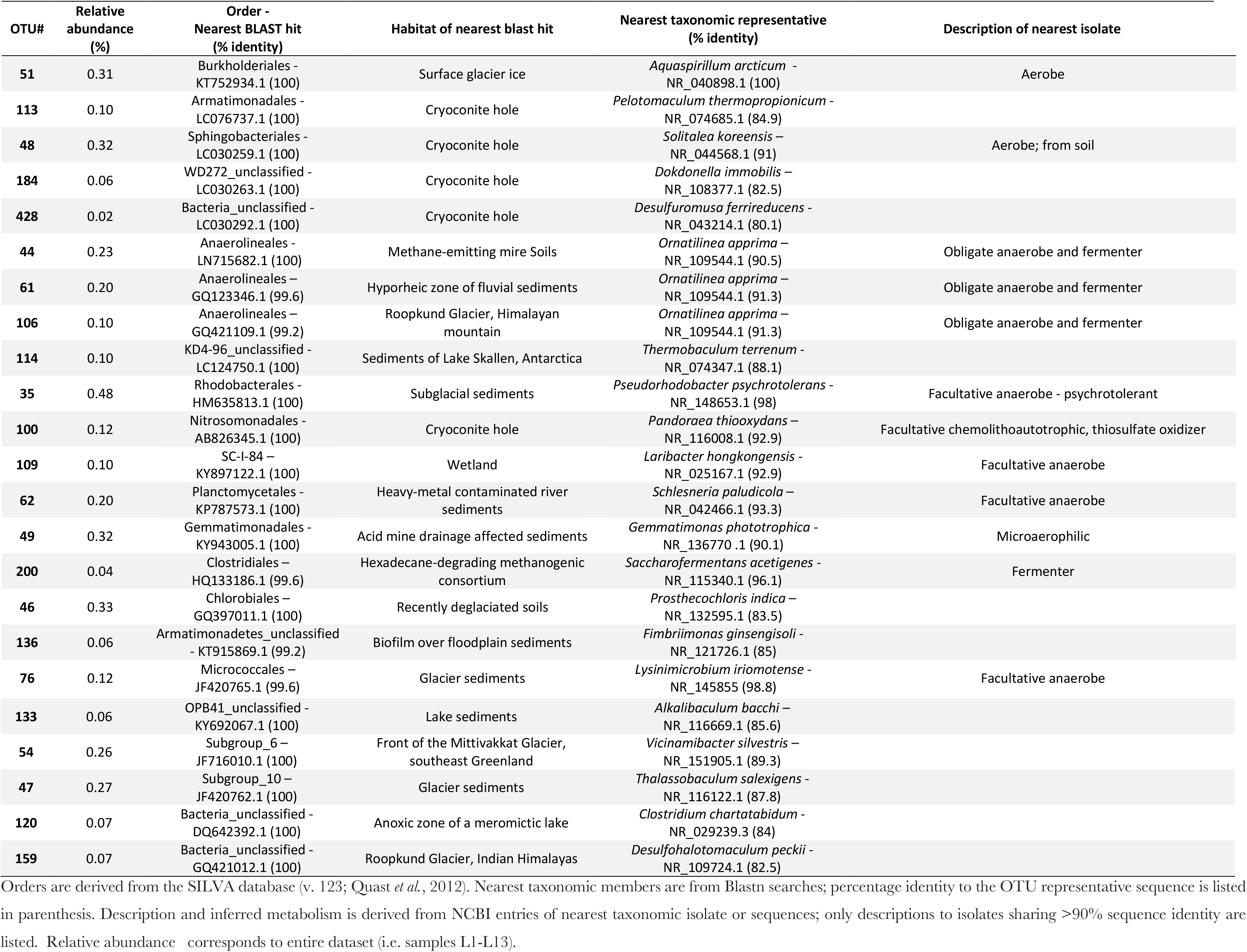
Taxonomic and inferred metabolic description of the representative sequence OTUs listed in Figure 4.

| OTU# | Relative abundance (%) | Order -<br>Nearest BLAST hit<br>(% identity) | Habitat of nearest blast hit | Nearest taxonomic representative<br>(% identity) | Description of nearest isolate |
| --- | --- | --- | --- | --- | --- |
| 51 | 0.31 | Burkholderiales -<br>KT752934.1 (100) | Surface glacier ice | *Aquaspirillum arcticum* -<br>NR_040898.1 (100) | Aerobe |
| 113 | 0.10 | Armatimonadales -<br>LC076737.1 (100) | Cryoconite hole | *Pelotomaculum thermopropionicum* -<br>NR_074685.1 (84.9) |  |
| 48 | 0.32 | Sphingobacteriales -<br>LC030259.1 (100) | Cryoconite hole | *Solitalea koreensis* -<br>NR_044568.1 (91) | Aerobe; from soil |
| 184 | 0.06 | WD272_unclassified -<br>LC030263.1 (100) | Cryoconite hole | *Dokdonella immobilis* -<br>NR_108377.1 (82.5) |  |
| 428 | 0.02 | Bacteria_unclassified -<br>LC030292.1 (100) | Cryoconite hole | *Desulfuromusa ferrireducens* -<br>NR_043214.1 (80.1) |  |
| 44 | 0.23 | Anaerolineales -<br>LN715682.1 (100) | Methane-emitting mire Soils | *Ornatilinea apprima* -<br>NR_109544.1 (90.5) | Obligate anaerobe and fermenter |
| 61 | 0.20 | Anaerolineales -<br>GQ123346.1 (99.6) | Hyporheic zone of fluvial sediments | *Ornatilinea apprima* -<br>NR_109544.1 (91.3) | Obligate anaerobe and fermenter |
| 106 | 0.10 | Anaerolineales -<br>GQ421109.1 (99.2) | Roopkund Glacier, Himalayan<br>mountain | *Ornatilinea apprima* -<br>NR_109544.1 (91.3) | Obligate anaerobe and fermenter |
| 114 | 0.10 | KD4-96_unclassified -<br>LC124750.1 (100) | Sediments of Lake Skallen, Antarctica | *Thermobaculum terrenum* -<br>NR_074347.1 (88.1) |  |
| 35 | 0.48 | Rhodobacterales -<br>HM635813.1 (100) | Subglacial sediments | *Pseudorhodobacter psychrotolerans* -<br>NR_148653.1 (98) | Facultative anaerobe - psychrotolerant |
| 100 | 0.12 | Nitrosomonadales -<br>AB826345.1 (100) | Cryoconite hole | *Pandoraea thiooxydans* -<br>NR_116008.1 (92.9) | Facultative chemolithoautotrophic, thiosulfate oxidizer |
| 109 | 0.10 | SC-I-84 -<br>KY897122.1 (100) | Wetland | *Laribacter hongkongensis* -<br>NR_025167.1 (92.9) | Facultative anaerobe |
| 62 | 0.20 | Planctomycetales -<br>KP787573.1 (100) | Heavy-metal contaminated river<br>sediments | *Schlesneria paludicola* -<br>NR_042466.1 (93.3) | Facultative anaerobe |
| 49 | 0.32 | Gemmatimonadales -<br>KY943005.1 (100) | Acid mine drainage affected sediments | *Gemmatimonas phototrophica* -<br>NR_136770.1 (90.1) | Microaerophilic |
| 200 | 0.04 | Clostridiales -<br>HQ133186.1 (99.6) | Hexadecane-degrading methanogenic<br>consortium | *Saccharofermentans acetigenes* -<br>NR_115340.1 (96.1) | Fermenter |
| 46 | 0.33 | Chlorobiales -<br>GQ397011.1 (100) | Recently deglaciated soils | *Prosthecochloris indica* -<br>NR_132595.1 (83.5) |  |
| 136 | 0.06 | Armatimonadetes_unclassified<br>- KT915869.1 (99.2) | Biofilm over floodplain sediments | *Fimbriimonas ginsengisoli* -<br>NR_121726.1 (85) |  |
| 76 | 0.12 | Micrococcales -<br>JF420765.1 (99.6) | Glacier sediments | *Lysinibaculum iriomotense* -<br>NR_145855 (98.8) | Facultative anaerobe |

|  |  |  |  |  |
| --- | --- | --- | --- | --- |
| 133 | 0.06 | OPB41_unclassified -<br>KY692067.1 (100) | Lake sediments | *Alkalibaculum bacchi* –<br>NR_116669.1 (85.6) |
| 54 | 0.26 | Subgroup_6 –<br>JF716010.1 (100) | Front of the Mittivakkat Glacier,<br>southeast Greenland | *Vicinamibacter silvestris* –<br>NR_151905.1 (89.3) |
| 47 | 0.27 | Subgroup_10 –<br>JF420762.1 (100) | Glacier sediments | *Thalassobaculum salexigens* -<br>NR_116122.1 (87.8) |
| 120 | 0.07 | Bacteria_unclassified -<br>DQ642392.1 (100) | Anoxic zone of a meromictic lake | *Clostridium chartatabidum* -<br>NR_029239.3 (84) |
| 159 | 0.07 | Bacteria_unclassified -<br>GQ421012.1 (100) | Roopkund Glacier, Indian Himalayas | *Desulfohalotomaculum peckii* -<br>NR_109724.1 (82.5) |

Orders are derived from the SILVA database (v. 123; Quast *et al.*, 2012). Nearest taxonomic members are from Blastn searches; percentage identity to the OTU representative sequence is listed in parenthesis. Description and inferred metabolism is derived from NCBI entries of nearest taxonomic isolate or sequences; only descriptions to isolates sharing >90% sequence identity are listed. Relative abundance corresponds to entire dataset (i.e. samples L1-L13).

## Discussion

The observation of a sustained core microbiome throughout the melt season at LG aligns with an increasing number of studies of proglacial meltwater systems (Dieser et al., 2014, Sheik et al., 2015, Kohler et al., 2020). Composition of exported communities during the 2015 melt-season also reflects that of a previous study in the region (Cameron et al., 2017), but also in glacial systems elsewhere (Kohler et al., 2020), and strengthens the view of major phylotypes characterising the subglacial margin of glaciers worldwide. For example, the dominance of Betaproteobacteria has previously been proposed to be a main constituent of subglacial communities (Boetius et al., 2015, Kohler et al., 2020), and Betaproteobacteria account for the majority of exported populations at LG (most Betaproteobacteria sequences fall within the order Burkholderiales at LG). The same is true for other major LG orders (Xanthomonadales, Sphingobacteriales, Micrococcales, etc.), which have also been shown to dominate previously monitored proglacial rivers (Kohler *et al.*, 2020).

Whether this conserved microbiome is representative of a relatively homogeneous, ‘blanket’ ecosystem beneath the ice, however, is difficult to reconcile from proglacial communities alone. The persistence of a core microbiota throughout the melt season (i.e. present in every sample) argues for some degree of homogenisation, either regarding the subglacial environment itself, or reflective of mixing processes during transport to the ice margin, or a combination of the two (e.g. Žárský *et al.*, 2018). As such, our view of community assemblages sampled at LG (and elsewhere) are likely biased towards more marginal populations, or those indigenous to flanking regions of main drainage channels beneath the ice (e.g. hyporheic-like regions; see Tranter *et al.* (2005)), “ironing-out” signals from more remote subglacial communities. That said, glacial hydrology has previously been demonstrated to exert (some degree of) control on microbial assemblages exported to the proglacial zone, and it is possible to link subglacial water residence-time to changes in community structures (Sheik et al., 2015, Dubnick et al., 2017). Here, the high-resolution hydrochemical and hydrological information from the proglacial river allows further inferences on the subglacial niche partitioning that may be present beneath larger ice-sheet catchments such as LG, which undergoes more complex hydrological evolution than its smaller counterparts (Davison et al., 2019).

### Hydrological forcing on exported microbial assemblages

Niche differentiation between a channelized and distributed system beneath the ice (Figs. 4) supports previously proposed conceptual models of subglacial habitats. Oxic meltwaters entering major drainage channels should maintain a more oxidizing environment capable of sustaining (micro)aerobic populations, and create a gradient of increasingly reducing/anoxic conditions with distance from main channels to distributed regions of the bed, more favourable to communities relying on more reduced electron acceptors (Tranter *et al.*, 2005, Hodson *et al.*, 2008). The relative decrease in abundance of putative aerobic methyl-an methanotrophic clades (e.g. OTUs, 3, 9, 16 Figs 3c; Table 1) at the expense of (putatively) anaerobic ones (e.g. of the order Anaerolineales; Fig. 3, 4; Tables 1, 2) during periods of enhanced subglacial flushing (outbursts) suggests such methylophilic populations may be more constrained to glacial/channel margins beneath the catchment, and agrees with previous models of increased hypoxia/anoxia away from main drainage channels. Anaerolineales also comprised a large fraction of phylotypes detected in small subglacial outflows of the neighbouring Russell Glacier, when supraglacially-sourced sequences were removed from bulk community observations (Dieser et al., 2014).

### Supraglacial runoff imprints on proglacial communities

OTUs significantly impacting community changes following the outburst period largely comprised of sequences related to those from glacier surfaces (e.g. cryoconite holes; Fig. 4c, Table 2), likely reflecting the dominance of rapid subglacial transport of lower residence-time waters via an efficient, channelised drainage system (Chandler *et al.*, 2013). This later-season trend at LG likely reflects that previously observed in smaller glacier systems that experience less dramatic hydrological change, not influenced by supraglacial lake-drainage (outburst) events. For example, Dubnick *et al.* (2017) described the evolution of the Kiattuut Sermiat drainage system as an ongoing dilution of the subglacial signal by supraglacial waters, with a microbiological signature of exported assemblages approaching that of supraglacial communities as the season progressed. Proglacial rivers fed by multiple glaciers on Qeqertasuaq (Disko Island, West Greenland) during the late melt-season in August 2015 also revealed large contributions of subglacial meltwaters to community assemblages (Žárský *et al.*, 2018).

### Colonisation of the ice sheet bed

Similarities between communities exported later in the season (L13) and those observed in front of the ice-sheet beneath the river ice in early May (L1) suggest some retention of late-season meltwater (populations) at the glacier’s margin and forefield, following hydrological shut-down at the end of the melt-season, and is consistent with recent study of glacier naled ice communities in Svalbard (Sułowicz *et al.*, 2020). This probably indicates that very early basal-melt waters carry a legacy of assemblages from the previous melt season, alongside populations derived from basal ice, and may explain the higher proportions of phylotypes related to glaciers’ surfaces (e.g. OTU 3) in L1 samples (Fig. 2, 4, Tables 1–2). It also indicates a stronger surface signal than previously expected might be retained subglacially, despite surface waters containing an order of magnitude less cells than those of the LG proglacial river (Cameron *et al.*, 2017). It should be noted that this “supraglacial legacy signature” inferred from microbial populations is “lost” when looking at water chemistry, which bears a strong basal signature typical of a distributed drainage system very early on in the melt season, and which is very distinct from that observed during the late season (Bartholomew *et al.*, 2011, Hindshaw *et al.*, 2014, Kellerman *et al.*, 2020). Late-season communities retained subglacially or at the glacier front likely undergo selection overwinter and therefore supraglacial influence is probably more important for generalist populations than those adapted to glacial surfaces such as *Cyanobacteria* (Gokul *et al.*, 2019). For example, *Cyanobacteria* here accounted for less than 0.03% of all sequences (data not shown), which at first glance would argue against a significant imprint of surface melt on the observed communities. The potential influence of glacier surface populations in shaping subglacial biota has recently been highlighted by a study of GrIS surface waters, which revealed a high-abundance of phylotypes related to those typically associated to subglacial systems (Gokul *et al.*, 2019).

### Inferred metabolic functions of exported microbial assemblages

Although we are aware of the limitations in unambiguously assigning metabolic functions to specific phylotypes based on (partial) 16S rRNA sequence information alone, comparison against the public repository can still inform on the putative metabolism of some LG populations. Amongst others, the representative sequence of major OTUs identically or closely matched those isolated from other cold and/or glaciated environments, as well as those involved in methane, sulphur, and iron cycling (Table1). The most abundant phylotype detected in all samples, OTU 1, perfectly matched partial 16S rRNA sequences of *Rhodoferax ferrireducens*, a psychrotolerant facultative anaerobe that can reduce Fe(III) using a range of simple organic compounds (Finneran *et al.*, 2003). *Rhodoferax* species have previously been shown to dominate iron-reducing enrichments with LG basal ice (Nixon *et al.*, 2016), making it likely that OTU 1 indeed carries on iron-reduction, and therefore that iron reduction plays a key function beneath the LG catchment. A strong potential for iron and sulphur (e.g. pyrite) oxidation is also highlighted by other dominant OTUs (OTUs 5, 14, and 20; Table 1), and allude to a complete iron cycle beneath the ice and its link to pyrite or other iron-sulphur cycling. As found in other glacial environments, these putative (iron)-sulphur oxidisers may also act as primary producers, supplying carbon to the subglacial system via chemolithoautotrophy (e.g. *Syderoxidans* – OTU 5, and *Thiobacillus* – OTU 14 *sp.;* Boyd *et al.* (2014)).

Methane cycling beneath LG has been previously demonstrated, as well as the presence of methanotrophs related to *Methylobacter tundripaludum* (OTU 9; Table 1), and to a lesser extent methanogens (Supp. Fig. tree/OTUs), in LG proglacial runoff (Lamarche-Gagnon *et al.*, 2019). Additional potential methanotrophic clades are further identified here (e.g. OTU 16; Fig). 16S rRNA results mostly agree with a recent metagenomic investigation of methane-cycling genes from the same sampling season (Rybár, 2020), re-enforcing the view on the function of putative methano-genic/trophic OTUs. That is, the temporal distribution of methanotrophic 16S rRNA sequences (elevated in early, “pre-melt” samples and post-outburst period, Fig. 3) was also reflected in a relative increase of *pmoA* genes (functional marker of aerobic methanotrophy) during the same overall periods (Rybár, 2020). Again similar to found with 16S rRNA sequencing, *mcrA* genes (functional marker for methanogenesis) were only recovered in very low quantities within the LG metagenomes, and were related to hydrogenotrophic clades (Rybár, 2020); most LG methanogen-related 16S rRNA sequences also belonged to known H2-oxidising taxa – as opposed to acetoclastic ones (Fig. S1-2; Lamarche-Gagnon *et al.*, 2019). A notable difference, however, was the absence of (*mcrA*) sequences related to anaerobic methane oxidising archaea (ANME) in the LG metagenomes (Rybár, 2020), which contrasts with 16S rRNA results here. The most abundant archaeal phylotype (OTU 137) beneath the river-ice in May and during the later season most closely relate to recently characterised anaerobic methane oxidisers of the clade ANME-2d, which couples methane oxidation to iron reduction (Fig. S2; Cai *et al.*, 2018). Similar sequences have also been identified in the sediments of the alpine Robertson Glacier, Canada, and inferred to belong to anaerobic methane oxidisers (Fig. S2; Boyd *et al.*, 2010). ANME-2d sequences have also been identified in anaerobic delta sediments of the Watson River, fed by LG meltwaters, and their methanotrophic activity suggested by long-term incubation experiments (Cameron *et al.*, 2017).

### A microbial window into the subglacial environment

The data presented here grant a glimpse through a microbial window into the LG subglacial system. We observe a subglacial microbiome potentially centred on chemo(auto)trophic iron-cycling (e.g. OTUs 1, 5, 20), supporting the idea that the high abundance of nanoparticulate iron (oxy)hydroxides found previously at LG are bioavailable and/or are the product of biogeochemical weathering (Hawkings *et al.*, 2014, Hawkings *et al.*, 2018). Methanotrophic and methylotrophic clades (e.g. OTUs 3, 9, 16) appear more prominent in well-aerated regions of the bed, or those seasonally influenced by supraglacial melt, including the proglacial zone all throughout the winter period (L1). We therefore find that supraglacial melt, likely the oxygen it carries, but also potentially allochthonous organic carbon and cells (Lawson *et al.*, 2014, Kellerman *et al.*, 2020), appears to shape the biota of the subglacial environment beneath the margin of the GrIS, as has been alluded to for small glacier systems (Tranter et al., 2005). The main drainage channels and ice margins seem to operate as a methanotrophic strip, relying on both oxygen supply from the surface and methane from deeper sediment/till layers, and more isolated sections of the bed (Lamarche-Gagnon et al., 2019). However, the very low abundance of methanogen phylotypes detected despite elevated concentrations of methane exported from the catchment (Lamarche-Gagnon et al., 2019) suggests that our snapshot is an incomplete portrayal of the LG subglacial ecosystem. Fermentation of old organic carbon may play a more important role in more remote sections of the bed given the relative increase of putative fermenters (e.g. of the order *Anaerolineales*; McIlroy *et al.*, 2017) exported during outburst events.

The ecosystem depicted here looks highly similar to that described in Whillans Subglacial Lake (SLW) beneath Antarctica, despite it being a far more isolated ice-sheet environment (e.g. not influenced by supraglacial melt) compared to the LG drainage system. There, the availability of methane, sulphur, iron, and oxygen (amongst other) also appears to shape microbial communities of the lake micro-oxic waters influenced by basal melt, and the more anoxic underlying lake sediments (Christner *et al.*, 2014, Michaud *et al.*, 2017). Interestingly, major SLW phylotypes were similar or identical (i.e. 100% sequence identity; data not shown) to those described here (i.e. *Rhodoferax* - there reported as *Albidiferax* - *Sideroxydans, Thiobacillus*, and *Methylobacter* species; Purcell *et al.*, 2014, Achberger *et al.*, 2016). The very low abundance of methanogens in SLW despite the high methane concentrations in sediments also resembles descriptions here. Some of the information obtained at LG can therefore likely be extended to a more general view of ice-sheet beds, further highlighting similarities between glacial environments worldwide (Kohler *et al.*, 2020), even under contrasting hydrological regimes. Still, a full picture of subglacial ecosystems may ultimately only be depicted via direct access to the subsurface through drilling operations, or potentially via more extensive sampling and sequencing efforts.

## Supporting information

mothur_logfiles

## Acknowledgment

We thank J. E. Hatton, A. D. Beaton, and A. J. Tedstone who assisted with fieldwork at LG. This research was supported by Czech Science Foundation grants (GACR; 15-17346Y and 18-12630S) to M.S. and a UK NERC grant (NE/J02399X/1) to A.M.A. for DNA analyses. This research was also part of the UK NERC funded DELVE programme (NERC grant NE/I008845/1 to J.L.W.). G.L.-G. was funded by the University of Bristol Scholarship Programme and a FRQNT Scholarship (number 185136). The work was also supported by a Leverhulme research fellowship to J.L.W. J.R.H. was supported by a European Union Horizon 2020 research and innovation grant under the Marie Sklodowska-Curie Actions fellowship ICICLES (grant agreement #793962). We also thank the Kangerlussuaq International Science Station, especially Rikka Møller, for support with field logistics.

## Notes

### Competing Interest Statement

The authors have declared no competing interest.

## References

Achberger AM, Christner BC, Michaud AB, et al. (2016) Microbial Community Structure of Subglacial Lake Whillans, West Antarctica. Frontiers in Microbiology 7.

Bartholomew I, Nienow P, Sole A, Mair D, Cowton T, Palmer S & Wadham J (2011) Supraglacial forcing of subglacial drainage in the ablation zone of the Greenland ice sheet. Geophysical Research Letters 38: n/a–n/a.

Boetius A, Anesio AM, Deming JW, Mikucki JA & Rapp JZ (2015) Microbial ecology of the cryosphere: sea ice and glacial habitats. Nature Reviews Microbiology.

Boyd ES, Skidmore M, Mitchell AC, Bakermans C & Peters JW (2010) Methanogenesis in subglacial sediments. Environ Microbiol Rep 2: 685–692.

Boyd ES, Hamilton TL, Havig JR, Skidmore ML & Shock EL (2014) Chemolithotrophic Primary Production in a Subglacial Ecosystem. Applied and environmental microbiology 80: 6146–6153.

Brown GH (2002) Glacier meltwater hydrochemistry. Applied Geochemistry 17: 855–883.

Burns R, Wynn PM, Barker P, et al. (2018) Direct isotopic evidence of biogenic methane production and efflux from beneath a temperate glacier. Scientific Reports 8: 17118.

Cai C, Leu AO, Xie G-J, Guo J, Feng Y, Zhao J-X, Tyson GW, Yuan Z & Hu S (2018) A methanotrophic archaeon couples anaerobic oxidation of methane to Fe(III) reduction. The ISME Journal 12: 1929–1939.

Cameron KA, Stibal M, Olsen NS, Mikkelsen AB, Elberling B & Jacobsen CS (2017) Potential Activity of Subglacial Microbiota Transported to Anoxic River Delta Sediments. Microbial Ecology 74: 6–9.

Cameron KA, Stibal M, Hawkings JR, Mikkelsen AB, Telling J, Kohler TJ, Gözdereliler E, Zarsky JD, Wadham JL & Jacobsen CS (2017) Meltwater export of prokaryotic cells from the Greenland ice sheet. Environmental Microbiology 19: 524–534.

Caporaso JG, Lauber CL, Walters WA, Berg-Lyons D, Lozupone CA, Turnbaugh PJ, Fierer N & Knight R (2011) Global patterns of 16S rRNA diversity at a depth of millions of sequences per sample. Proceedings of the National Academy of Sciences 108: 4516–4522.

Chandler DM, Wadham JL, Lis GP, et al. (2013) Evolution of the subglacial drainage system beneath the Greenland Ice Sheet revealed by tracers. Nature Geosci 6: 195–198.

Christiansen JR & Jørgensen CJ (2018) First observation of direct methane emission to the atmosphere from the subglacial domain of the Greenland Ice Sheet. Scientific Reports 8: 16623.

Christner BC, Priscu JC, Achberger AM, et al. (2014) A microbial ecosystem beneath the West Antarctic ice sheet. Nature 512: 310–313.

Clason CC, Mair DWF, Nienow PW, Bartholomew ID, Sole A, Palmer S & Schwanghart W (2015) Modelling the transfer of supraglacial meltwater to the bed of Leverett Glacier, Southwest Greenland. The Cryosphere 9: 123–138.

Cowton T, Nienow P, Bartholomew I, Sole A & Mair D (2012) Rapid erosion beneath the Greenland ice sheet. Geology 40: 343–346.

Davison BJ, Sole AJ, Livingstone SJ, Cowton TR & Nienow PW (2019) The Influence of Hydrology on the Dynamics of Land-Terminating Sectors of the Greenland Ice Sheet. Frontiers in Earth Science 7.

Dawes PR (2009) The bedrock geology under the Inland Ice: the next major challenge for Greenland mapping. Geological Survey of Denmark and Greenland Bulletin 17: 57–60.

Dieser M, Broemsen EL, Cameron KA, King GM, Achberger A, Choquette K, Hagedorn B, Sletten R, Junge K & Christner BC (2014) Molecular and biogeochemical evidence for methane cycling beneath the western margin of the Greenland Ice Sheet. ISME J 8: 2305–2316.

Doyle SM, Montross SN, Skidmore ML & Christner BC (2013) Characterizing microbial diversity and the potential for metabolic function at-15° C in the Basal Ice of Taylor Glacier, Antarctica. Biology 2: 1034–1053.

Dubnick A, Kazemi S, Sharp M, Wadham J, Hawkings J, Beaton A & Lanoil B (2017) Hydrological controls on glacially exported microbial assemblages. Journal of Geophysical Research: Biogeosciences 122: 1049–1061.

Finneran KT, Johnsen CV & Lovley DR (2003) Rhodoferax ferrireducens sp. nov., a psychrotolerant, facultatively anaerobic bacterium that oxidizes acetate with the reduction of Fe (III). International Journal of Systematic and Evolutionary Microbiology 53: 669–673.

Gokul JK, Cameron KA, Irvine-Fynn TDL, Cook JM, Hubbard A, Stibal M, Hegarty M, Mur LAJ & Edwards A (2019) Illuminating the dynamic rare biosphere of the Greenland Ice Sheet’s Dark Zone. FEMS Microbiology Ecology 95.

Hatton JE, Hendry KR, Hawkings JR, Wadham JL, Kohler TJ, Stibal M, Beaton AD, Bagshaw EA & Telling J (2019) Investigation of subglacial weathering under the Greenland Ice Sheet using silicon isotopes. Geochimica et Cosmochimica Acta.

Hawkings JR, Wadham JL, Benning LG, Hendry KR, Tranter M, Tedstone A, Nienow P & Raiswell R (2017) Ice sheets as a missing source of silica to the polar oceans. Nature Communications 8: 14198.

Hawkings JR, Wadham JL, Tranter M, Raiswell R, Benning LG, Statham PJ, Tedstone A, Nienow P, Lee K & Telling J (2014) Ice sheets as a significant source of highly reactive nanoparticulate iron to the oceans. Nat Commun 5: 3929.

Hawkings JR, Benning LG, Raiswell R, Kaulich B, Araki T, Abyaneh M, Stockdale A, Koch-Müller M, Wadham JL & Tranter M (2018) Biolabile ferrous iron bearing nanoparticles in glacial sediments. Earth and Planetary Science Letters 493: 92–101.

Hawkings JR, Wadham JL, Tranter M, et al. (2015) The effect of warming climate on nutrient and solute export from the Greenland Ice Sheet. Geochemical Perspectives Letters 1: 94–104.

Hindshaw RS, Rickli J, Leuthold J, Wadham J & Bourdon B (2014) Identifying weathering sources and processes in an outlet glacier of the Greenland Ice Sheet using Ca and Sr isotope ratios. Geochimica et Cosmochimica Acta 145: 50–71.

Hodson A, Anesio AM, Tranter M, Fountain A, Osborn M, Priscu J, Laybourn-Parry J & Sattler B (2008) Glacial ecosystems. Ecological Monographs 78: 41–67.

Kellerman AM, Hawkings JR, Wadham JL, Kohler TJ, Stibal M, Grater E, Marshall M, Hatton JE, Beaton A & Spencer RGM (2020) Glacier Outflow Dissolved Organic Matter as a Window Into Seasonally Changing Carbon Sources: Leverett Glacier, Greenland. Journal of Geophysical Research: Biogeosciences 125: e2019JG005161.

Kohler TJ, Žárský JD, Yde JC, Lamarche-Gagnon G, Hawkings JR, Tedstone AJ, Wadham JL, Box JE, Beaton AD & Stibal M (2017) Carbon dating reveals a seasonal progression in the source of particulate organic carbon exported from the Greenland Ice Sheet. Geophysical Research Letters 6209–6217.

Kohler TJ, Vinšová P, Falteisek L, et al. (2020) Patterns in Microbial Assemblages Exported From the Meltwater of Arctic and Sub-Arctic Glaciers. Frontiers in Microbiology 11.

Kozich JJ, Westcott SL, Baxter NT, Highlander SK & Schloss PD (2013) Development of a Dual-Index Sequencing Strategy and Curation Pipeline for Analyzing Amplicon Sequence Data on the MiSeq Illumina Sequencing Platform. Applied and environmental microbiology 79: 5112–5120.

Lamarche-Gagnon G, Wadham JL, Sherwood Lollar B, et al. (2019) Greenland melt drives continuous export of methane from the ice-sheet bed. Nature 565: 73–77.

Lawson EC, Wadham JL, Tranter M, Stibal M, Lis GP, Butler CEH, Laybourn-Parry J, Nienow P, Chandler D & Dewsbury P (2014) Greenland Ice Sheet exports labile organic carbon to the Arctic oceans. Biogeosciences 11: 4015–4028.

McIlroy SJ, Kirkegaard RH, Dueholm MS, Fernando E, Karst SM, Albertsen M & Nielsen PH (2017) Culture-Independent Analyses Reveal Novel Anaerolineaceae as Abundant Primary Fermenters in Anaerobic Digesters Treating Waste Activated Sludge. Frontiers in Microbiology 8.

McMurdie P & Holmes S (2013) phyloseq: An R Package for Reproducible Interactive Analysis and Graphics of Microbiome Census Data. PLoS ONE 8: e61217–e61217.

Michaud AB, Dore JE, Achberger AM, Christner BC, Mitchell AC, Skidmore ML, Vick-Majors TJ & Priscu JC (2017) Microbial oxidation as a methane sink beneath the West Antarctic Ice Sheet. Nature Geosci 10: 582–586.

Mitchell AC, Lafreniere MJ, Skidmore ML & Boyd ES (2013) Influence of bedrock mineral composition on microbial diversity in a subglacial environment. Geology 41: 855–858.

Montross S, Skidmore M, Christner B, Samyn D, Tison J-L, Lorrain R, Doyle S & Fitzsimons S (2013) Debris-Rich Basal Ice as a Microbial Habitat, Taylor Glacier, Antarctica. Geomicrobiology Journal 31: 76–81.

Montross SN, Skidmore M, Tranter M, Kivimäki A-L & Parkes RJ (2013) A microbial driver of chemical weathering in glaciated systems. Geology 41: 215–218.

Nixon SL, Telling J, Wadham JL & Cockell CS (2016) Viable cold-tolerant iron-reducing microorganisms in geographically-isolated subglacial environments. Biogeosciences Discuss 2016: 1–19.

Oksanen J, Blanchet F, Friendly M, Kindt R, Legendre P & McGlinn D (2019) vegan: Community Ecology Package. R package version 2.5-6. 2019. p.^pp. https://CRAN.R-project.org/package=vegan.

Palmer S, Shepherd A, Nienow P & Joughin I (2011) Seasonal speedup of the Greenland Ice Sheet linked to routing of surface water. Earth and Planetary Science Letters 302: 423–428.

Purcell AM, Mikucki JA, Achberger A, et al. (2014) Microbial sulfur transformations in sediments from Subglacial Lake Whillans. Frontiers in Microbiology 5.

Quast C, Pruesse E, Gerken J, Peplies J, Yarza P, Yilmaz P, Schweer T & Glöckner FO (2012) The SILVA ribosomal RNA gene database project: improved data processing and web-based tools. Nucleic Acids Research 41: D590–D596.

Rybár M (2020) Genetic potential for methane metabolism in the Greenland subglacial ecosystem. MSc Thesis, Charles University.

Schloss PD, Westcott SL, Ryabin T, et al. (2009) Introducing mothur: Open-Source, Platform-Independent, Community-Supported Software for Describing and Comparing Microbial Communities. Applied and environmental microbiology 75: 7537–7541.

Sharp M, Parkes J, Cragg B, Fairchild IJ, Lamb H & Tranter M (1999) Widespread bacterial populations at glacier beds and their relationship to rock weathering and carbon cycling. Geology 27: 107–110.

Sheik CS, Stevenson EI, Den Uyl PA, Arendt CA, Aciego SM & Dick GJ (2015) Microbial communities of the Lemon Creek Glacier show subtle structural variation yet stable phylogenetic composition over space and time. Frontiers in Microbiology 6.

Skidmore M, Anderson SP, Sharp M, Foght J & Lanoil BD (2005) Comparison of Microbial Community Compositions of Two Subglacial Environments Reveals a Possible Role for Microbes in Chemical Weathering Processes. Applied and environmental microbiology 71: 6986–6997.

Skidmore ML, Foght JM & Sharp MJ (2000) Microbial Life beneath a High Arctic Glacier. Applied and environmental microbiology 66: 3214–3220.

Stibal M, Hasan F, Wadham JL, Sharp MJ & Anesio AM (2012) Prokaryotic diversity in sediments beneath two polar glaciers with contrasting organic carbon substrates. Extremophiles 16: 255–265.

Stibal M, Wadham JL, Lis GP, et al. (2012) Methanogenic potential of Arctic and Antarctic subglacial environments with contrasting organic carbon sources. Global Change Biology 18: 3332–3345.

Sułowicz S, Bondarczuk K, Ignatiuk D, Jania JA & Piotrowska-Seget Z (2020) Microbial communities from subglacial water of naled ice bodies in the forefield of Werenskioldbreen, Svalbard. Science of The Total Environment 723: 138025.

Team RC (2018) R: A language and environment for statistical computing. R Foundation for Statistical Computing. p.^pp.

Tranter M, Skidmore M & Wadham J (2005) Hydrological controls on microbial communities in subglacial environments. Hydrological Processes 19: 995–998.

Tranter M, Sharp MJ, Lamb HR, Brown GH, Hubbard BP & Willis IC (2002) Geochemical weathering at the bed of Haut Glacier d’Arolla, Switzerland—a new model. Hydrological Processes 16: 959–993.

Wadham JL, Tranter M, Skidmore M, Hodson AJ, Priscu J, Lyons WB, Sharp M, Wynn P & Jackson M (2010) Biogeochemical weathering under ice: Size matters. Global Biogeochem Cycles 24: n/a–n/a.

Wadham JL, Arndt S, Tulaczyk S, et al. (2012) Potential methane reservoirs beneath Antarctica. Nature 488: 633–637.

Warnes GR, Bolker B, Bonebakker L, Gentleman R, Liaw WHA, Lumley T, Maechler M, Magnusson A, Moeller S & Schwartz M (2019) gplots: Various R programming tools for plotting data. p.^pp. https://CRAN.R-project.org/package=gplots.

Wilhelm L, Singer GA, Fasching C, Battin TJ & Besemer K (2013) Microbial biodiversity in glacier-fed streams. ISME J 7: 1651–1660.

Yde JC, Finster KW, Raiswell R, Steffensen JP, Heinemeier J, Olsen J, Gunnlaugsson HP & Nielsen OB (2010) Basal ice microbiology at the margin of the Greenland ice sheet. Annals of Glaciology 51: 71–79.

Žárský JD, Kohler TJ, Yde JC, Falteisek L, Lamarche-Gagnon G, Hawkings JR, Hatton JE & Stibal M (2018) Prokaryotic assemblages in suspended and subglacial sediments within a glacierized catchment on Qeqertarsuaq (Disko Island), west Greenland. FEMS Microbiology Ecology 94: fiy100–fiy100.

