## Supplementary Information for "Meltwater runoff from the Greenland Ice Sheet reveals microbial consortia from contrasting subglacial drainage systems"

**Table S1.** Sampling and grouping details of collected Sterivex filter samples

| Group | Sampling Time | Sterivex code | Replicate label | Volume (mL) | Sampling Site |
|---|---|---|---|---|---|
| L1 | 2015-05-04 18:00 | MP14 | 1A | 6000 | Portal/borehole |
| L1 | 2015-05-04 20:00 | L2 | 1A | 1800 | Portal/borehole |
| L1 | 2015-05-10 11:00 | MP16 | 1B | 5000 | Portal/chainsaw |
| L1 | 2015-05-13 12:00 | MP13 | 1C | 4500 | Portal/chainsaw |
| L3 | 2015-06-07 12:00 | L1 | 3A | 400 | Upwelling |
| L3 | 2015-06-07 12:00 | T2 | 3B | 400 | Upwelling |
| L4 | 2015-06-20 12:30 | M1 | 4A | 600 | Cliff |
| L4 | 2015-06-20 17:00 | MP1 | 4B | 600 | Cliff |
| L4 | 2015-06-20 17:00 | W1 | 4C | 600 | Cliff |
| L4 | 2015-06-20 18:00 | L1* | 4ABC | 500 | Cliff |
| L4 | 2015-06-20 16:00 | W6 | 4ABC | 1000 | Cliff portal side |
| L5 | 2015-06-23 16:00 | MP2 | 5A | 600 | Cliff |
| L5 | 2015-06-23 16:00 | MP3 | 5B | 600 | Cliff |
| L5 | 2015-06-23 16:00 | MP4 | 5C | 540 | Cliff |
| L6 | 2015-06-25 17:00 | MP5 | 6A | 600 | Cliff |
| L6 | 2015-06-25 17:00 | MP6 | 6B | 600 | Cliff |
| L6 | 2015-06-25 17:00 | W2 | 6C | 600 | Cliff |
| L6 | 2015-06-25 17:00 | L2* | 6ABC | 600 | Cliff |
| L6 | 2015-06-27 19:00 | MP7 | 6A | 600 | Cliff |
| L6 | 2015-06-27 19:00 | MP8 | 6B | 600 | Cliff |
| L6 | 2015-06-27 19:00 | MP9 | 6C | 600 | Cliff |
| L6 | 2015-06-27 19:00 | W4 | 6ABC | 700 | Cliff |
| L8 | 2015-07-02 12:00 | M2 | 8A | 600 | Cliff |
| L8 | 2015-07-02 12:00 | M3 | 8B | 600 | Cliff |
| L8 | 2015-07-02 12:00 | M4 | 8C | 600 | Cliff |
| L9 | 2015-07-05 12:00 | M5 | 9A | 600 | Cliff |
| L9 | 2015-07-05 12:00 | M6 | 9B | 600 | Cliff |
| L9 | 2015-07-05 12:00 | M7 | 9C | 600 | Cliff |
| L9 | 2015-07-05 12:00 | W3 | 9ABC | 600 | Cliff |
| L9 | 2015-07-05 12:00 | W5 | 9ABC | 600 | Cliff |
| L10 | 2015-07-07 10:00 | T3 | 10A | 740 | Cliff |
| L10 | 2015-07-07 10:00 | T4 | 10B | 750 | Cliff |
| L10 | 2015-07-07 10:00 | T5 | 10ABC | 720 | Cliff |
| L10 | 2015-07-07 10:00 | T6 | 10C | 750 | Cliff |
| L11 | 2015-07-10 11:30 | M8 | 11A | 600 | Cliff |
| L11 | 2015-07-10 11:30 | M9 | 11B | 600 | Cliff |
| L11 | 2015-07-10 11:30 | M10 | 11C | 600 | Cliff |
| L12 | 2015-07-13 11:30 | MP10 | 12A | 600 | Cliff |
| L12 | 2015-07-13 11:30 | MP11 | 12B | 600 | Cliff |
| L12 | 2015-07-13 11:30 | MP12 | 12C | 600 | Cliff |
| L12 | 2015-07-17 00:00 | M11 | 12A | 600 | Cliff |
| L12 | 2015-07-17 00:00 | M12 | 12B | 600 | Cliff |
| L12 | 2015-07-17 00:00 | M13 | 12C | 600 | Cliff |
| L13 | 2015-07-21 12:00 | LA | 13A | 600 | Cliff |
| L13 | 2015-07-21 12:00 | LB | 13B | 600 | Cliff |
| L13 | 2015-07-21 12:00 | LC | 13C | 600 | Cliff |
| L13 | 2015-07-26 11:00 | M14 | 13A | 600 | Cliff |
| L13 | 2015-07-26 11:00 | M15 | 13B | 600 | Cliff |
| L13 | 2015-07-26 11:00 | M16 | 13C | 600 | Cliff |

“Sterivex code” refers to individual Sterivex samples prior to pooling into replicates; codes refer to “in-house” archiving. Volume refers to the approximate volume of filtrate. The portal borehole, chainsaw, and upwelling sites are described in Lamarche-Gagnon et al. (2019); the “cliff” site is the same as described in the supplementary material of Kohler *et al.* (2017) (Fig. 1).

mothur commands; batch file

mothur was run in “batch mode” on a remote server. The complete mothur “logfile” is included as supplementary information. Initial steps deviated from the mothur MiSeq standard operation procedure (Kozich *et al.*, 2013). Paired sequences from the sequencing centre contained both “forward” and “reverse” reads in both “R1” and “R2” fastq files, preventing the generation of a “.files” or “.oligos” file in mothur. Instead, the command “trim.seqs” with an “.oligos” file was used after converting raw reads into contigs. The batch file used to run initial mothur commands for quality checks and OTU generation is also included as supplementary information.

Table S2 summarises the number of reads in each samples following the “trim.seqs” (initial stage) and “remove.lineage” (final stage) commands.

**Table S2**

| SampleID | Initial | Final |
|---|---|---|
| L10A | 90955 | 78855 |
| L10B | 100653 | 87914 |
| L10C | 78594 | 69407 |
| L11A | 97354 | 85803 |
| L11B | 102383 | 89316 |
| L11C | 120466 | 105580 |
| L12A | 127498 | 110751 |
| L12B | 103948 | 90591 |
| L12C | 112485 | 98392 |
| L13A | 96380 | 83611 |
| L13B | 98034 | 82311 |
| L13C | 109065 | 89166 |
| L1A | 93389 | 69540 |
| L1B | 129635 | 102590 |
| L1C | 128344 | 99971 |
| L3A | 134685 | 112431 |
| L3B | 122893 | 103475 |
| L4A | 111038 | 95471 |
| L4B | 102506 | 87982 |
| L4C | 90831 | 78985 |
| L5A | 114118 | 98126 |
| L5B | 79782 | 69311 |

| | | |
|---|---|---|
| L5C | 104764 | 90167 |
| L6A | 109858 | 94172 |
| L6B | 70815 | 62718 |
| L6C | 88473 | 78229 |
| **L7A** | **41647** | **36859** |
| **L7B** | **30828** | **28083** |
| **L7C** | **45686** | **41114** |
| **L8A** | **37069** | **33253** |
| L8B | 95257 | 81292 |
| L8C | 99085 | 85050 |
| L9A | 118410 | 100998 |
| L9B | 121614 | 105199 |
| L9C | 108365 | 92715 |
| Total (unique) | - | 271165 |
| Total | 3416907 | 2919428 |

Highlighted in bold are samples with a lower number of reads that were excluded from downstream analyses as a result of subsampling (see above).

"Total (unique)" refers to the total number of reads that did not share 100% sequence identity; i.e. were "unique".

mothur commands; local machine

Commands in mothur used to remove doubletons (i.e. OTUs containing 2 sequences or less amongst all samples), subsample groups to the same level of sequences, calculate a Bray-Curtis (dis)similarity matrix, and generate ordinations were performed in mothur v.1.37.5 on a local machine. These commands respectively were "remove.rare(shared=Lev.shared, nseqs=2, label=0.03)", "sub.sample(shared=current, size=62242)", which eliminated samples L7s and L8A, and "dist.shared(shared=current, calc=braycurtis, label=0.03)". PCoA and nMDS ordinations were calculated on the generated Bray-Curtis matrix using the "pcoa" and "nmds" commands in mothur, respectively. The mothur outputted mothur logfile for the commands performed on the local machine (laptop) is also included as supplementary information.

**Table S3.** Major ion and dissolved silica (DSi) concentrations as well as hydrochemical sensor measurements for borehole and chainsaw hole samples

| Location | Sampling time | Julian Day | DSi | $F^-$ | $Cl^-$ | $SO_4^{2-}$ | $NO_3^-$ | $Na^+$ | $K^+$ | $Mg^{2+}$ | $Ca^{2+}$ | $HCO_3^-$ | D:M | $O_2$ (μM) | pH | EC (μS cm-1) |
|---|---|---|---|---|---|---|---|---|---|---|---|---|---|---|---|---|
| **Borehole** | 04/05/2015 17:00 | 124.7 | 164 | 8 | 54 | 513 | 0.52 | 98 | 76 | 222 | 978 | 800 | 6.9 | | | |
| **Borehole** | 04/05/2015 17:00 | 124.7 | 158 | 6.6 | 28 | 513 | 0.36 | 99 | 76 | 220 | 836 | 801 | 6.0 | | | |
| **Borehole** | 04/05/2015 17:00 | 124.7 | 164 | 7.1 | 29 | 511 | 0.34 | 99 | 76 | 224 | 846 | 802 | 6.1 | 25.8 (0.4) | 8.0 (0.2) | 70.7 (1.7) |
| **Borehole** | 04/05/2015 18:00 | 124.8 | 163 | 3.8 | 21 | 512 | 0.46 | 98 | 76 | 223 | 851 | 711 | 6.2 | | | |
| **Borehole** | 04/05/2015 20:00 | 124.8 | 164 | 4.5 | 22 | 514 | 0.46 | 99 | 77 | 223 | 855 | 713 | 6.1 | | | |
| **Chainsaw hole** | 10/05/2015 13:15 | 130.6 | 156 | 4.9 | 28 | 521 | 0.70 | 101 | 78 | 225 | 852 | 700 | 6.0 | ND | ND | ND |
| **Chainsaw hole** | 13/05/2015 12:00 | 133.5 | 148 | 6.5 | 28 | 476 | 1.61 | 93 | 71 | 207 | 775 | 634 | 6.0 | ND | ND | ND |

Major ion and DSi concentrations are reported as μeq $L^{-1}$ and μM, respectively. Values for sensor measurements ($O_2$, pH and EC) are averages from multiple deployments from 19:40 to 20:40 in depths ranging from ~ 2-5 meters below the river-ice surface. The number of measurements for O2, pH and EC respectively are 500, 39, and 16, with standard deviations reported in parenthesis. ND refers to "not-determined".

### Supplementary Figures

**Figure S1.** Relative abundance of major archaeal OTUs in LG microbial assemblages. Data is shown as both box plot (left) and bar plot (right); only the top 10 taxa are shown. The box mid-lines represent medians; the interquartile range (IQR) is represented by the lower and upper box boundaries, which denote the 25th and 75th percentiles, respectively; whiskers indicate confidence intervals 1.5 times the IQR, and points are outliers. Colour of points correspond to sampling time – same colour scheme as Figure 2 is used. Taxonomic information down to the order level (separated by dashes) for each OTU, when available, is indicated in the barplot legend.

**Figure S2.** Maximum likelihood trees of archaeal 16S rRNA gene sequences related to methanogens and ANME clades. This tree is an adaption of the one presented in Lamarche-Gagnon *et al.* (2019). Tree is rooted with the sequences of *Acidibilus sulfurireducens* and *Caldisphaera draconis*. LG OTUs are highlighted in bold with coloured squares identical to those found in the bar plot of Fig. S1 for OTUs (OTU 1115 is not shown on Fig. S1). ANME2-d related sequences recovered from Roberston Glacier (Boyd *et al.*, 2010) are also highlighted in bold. Only bootstrap vales > 50% are shown on tree nodes.

### References


Boyd ES, Skidmore M, Mitchell AC, Bakermans C & Peters JW (2010) Methanogenesis in subglacial sediments. *Environ Microbiol Rep* **2**: 685-692.

Kohler TJ, Žárský JD, Yde JC, Lamarche-Gagnon G, Hawkings JR, Tedstone AJ, Wadham JL, Box JE, Beaton AD & Stibal M (2017) Carbon dating reveals a seasonal progression in the source of particulate organic carbon exported from the Greenland Ice Sheet. *Geophysical Research Letters* 6209-6217.

Kozich JJ, Westcott SL, Baxter NT, Highlander SK & Schloss PD (2013) Development of a Dual-Index Sequencing Strategy and Curation Pipeline for Analyzing Amplicon Sequence Data on the MiSeq Illumina Sequencing Platform. *Applied and environmental microbiology* **79**: 5112-5120.

Lamarche-Gagnon G, Wadham JL, Sherwood Lollar B, *et al.* (2019) Greenland melt drives continuous export of methane from the ice-sheet bed. *Nature* **565**: 73-77.

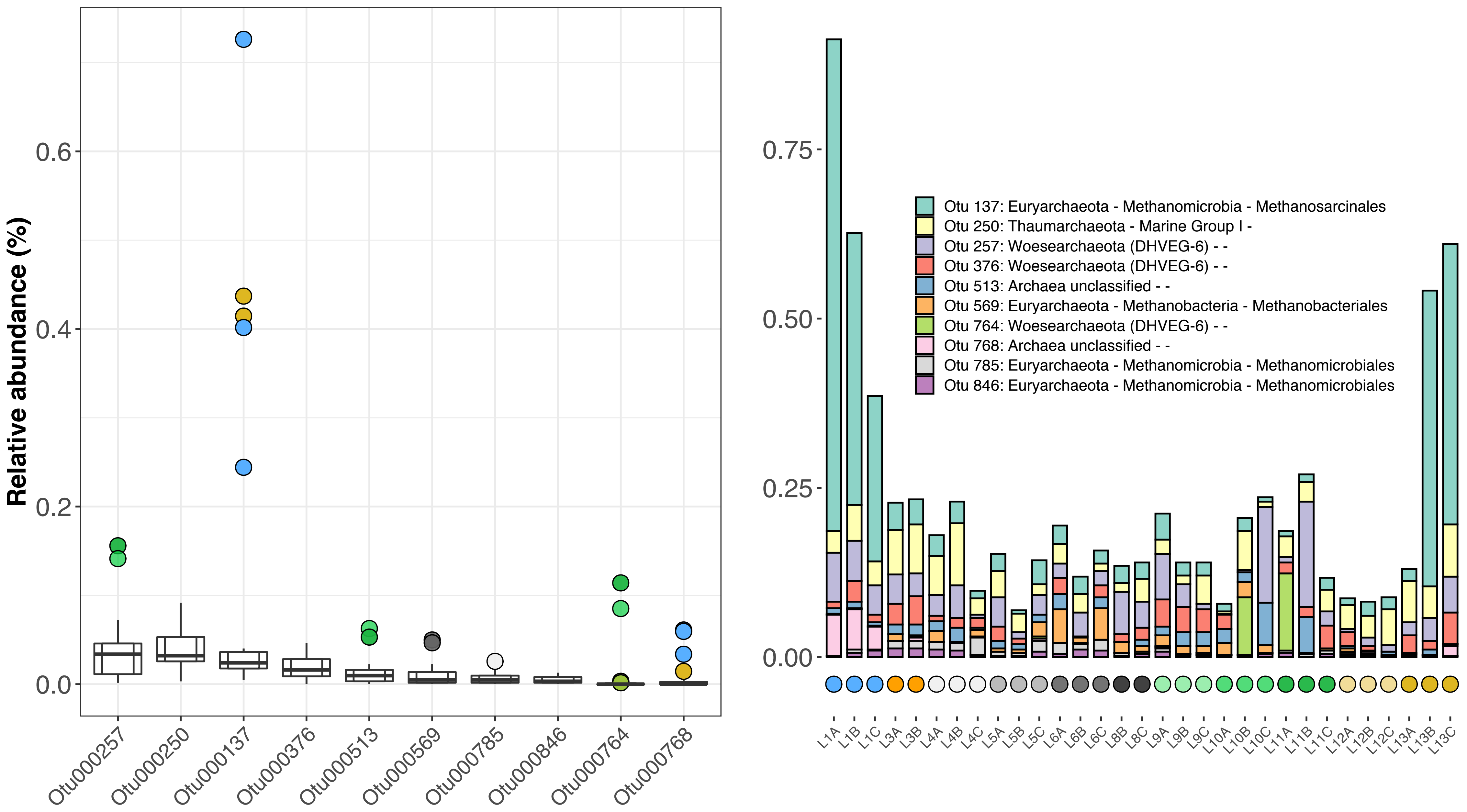

Relative abundance (%)
0.0
0.2
0.4
0.6
Otu000257
Otu000250
Otu000137
Otu000376
Otu000513
Otu000569
Otu000785
Otu000846
Otu000764
Otu000768
0.00
0.25
0.50
0.75
Otu 137: Euryarchaeota - Methanomicrobia - Methanosarcinales
Otu 250: Thaumarchaeota - Marine Group I -
Otu 257: Woesearchaeota (DHVEG-6) - -
Otu 376: Woesearchaeota (DHVEG-6) - -
Otu 513: Archaea unclassified - -
Otu 569: Euryarchaeota - Methanobacteria - Methanobacteriales
Otu 764: Woesearchaeota (DHVEG-6) - -
Otu 768: Archaea unclassified - -
Otu 785: Euryarchaeota - Methanomicrobia - Methanomicrobiales
Otu 846: Euryarchaeota - Methanomicrobia - Methanomicrobiales
L1A L1B L1C L3A L3B L4A L4B L4C L5A L5B L5C L6A L6B L6C L8B L8C L9A L9B L9C L10A L10B L10C L11A L11B L11C L12A L12B L12C L13A L13B L13C

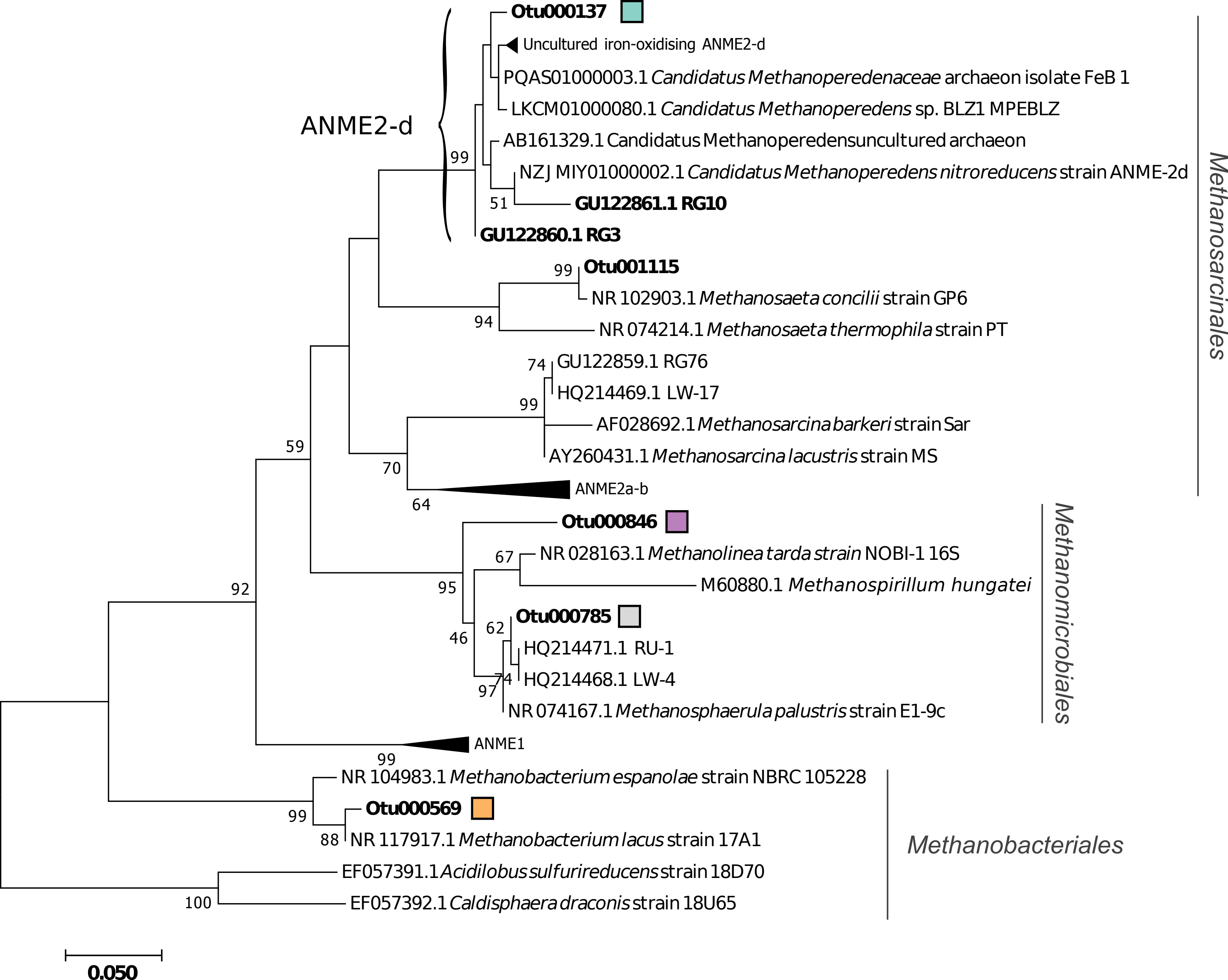
Otu000137
Uncultured iron-oxidising ANME2-d
PQAS01000003.1 Candidatus Methanoperedenaceae archaeon isolate FeB 1
LKCM01000080.1 Candidatus Methanoperedens sp. BLZ1 MPEBLZ
ANME2-d
AB161329.1 Candidatus Methanoperedensuncultured archaeon
99
NZJ MIY01000002.1 Candidatus Methanoperedens nitroreducens strain ANME-2d
51
GU122861.1 RG10
GU122860.1 RG3
99
Otu001115
NR 102903.1 Methanosaeta concilii strain GP6
94
NR 074214.1 Methanosaeta thermophila strain PT
74
GU122859.1 RG76
HQ214469.1 LW-17
99
AF028692.1 Methanosarcina barkeri strain Sar
59
AY260431.1 Methanosarcina lacustris strain MS
70
ANME2a-b
64
Methanosarcinales
Otu000846
67
NR 028163.1 Methanolinea tarda strain NOBI-1 16S
92
95
M60880.1 Methanospirillum hungatei
Otu000785
62
46
HQ214471.1 RU-1
74
HQ214468.1 LW-4
97
NR 074167.1 Methanosphaerula palustris strain E1-9c
Methanomicrobiales
ANME1
99
NR 104983.1 Methanobacterium espanolae strain NBRC 105228
Otu000569
99
88
NR 117917.1 Methanobacterium lacus strain 17A1
Methanobacteriales
EF057391.1 Acidilobus sulfurireducens strain 18D70
100
EF057392.1 Caldisphaera draconis strain 18U65
0.050
